# Opi1-mediated transcriptional modulation orchestrates genotoxic stress response in budding yeast

**DOI:** 10.1101/2022.11.04.515212

**Authors:** Giovanna Marques Panessa, Eduardo Tassoni-Tsuchida, Marina Rodrigues Pires, Rodrigo Rodrigues Felix, Rafaella Jekabson, Nadja Cristhina de Souza-Pinto, Fernanda Marques da Cunha, Onn Brandman, José Renato Rosa Cussiol

## Abstract

In budding yeast, the transcriptional repressor Opi1 regulates phospholipid biosynthesis by repressing expression of genes containing inositol-sensitive upstream activation sequences (UAS_INO_). Upon genotoxic stress, cells activate the DNA Damage Response (DDR) to coordinate a complex network of signaling pathways aimed at preserving genomic integrity. Here, we reveal that Opi1 is important to modulate transcription in response to genotoxic stress. We find that cells lacking Opi1 exhibit hypersensitivity to genotoxins, along with a delayed G1 to S-phase transition and decreased gamma-H2A levels. Transcriptome analysis using RNA-seq reveals that Opi1 plays a central role in modulating essential biological processes during genotoxic stress induced by methyl methanesulfonate, including repression of phospholipid biosynthesis and transduction of mating signaling. Moreover, Opi1 induces sulfate assimilation and amino acid metabolic processes, such as arginine and histidine biosynthesis and glycine catabolism. Furthermore, we observe increased mitochondrial DNA instability in *opi1Δ* cells upon MMS treatment. Notably, we show that constitutive activation of the transcription factors Ino2-Ino4 is responsible for genotoxin sensitivity in Opi1-deficient cells, and the production of inositol pyrophosphates by Kcs1 counteracts Opi1 function specifically during MMS-induced genotoxic stress. Overall, our findings highlight Opi1 as a critical sensor of genotoxic stress in budding yeast, orchestrating gene expression to facilitate appropriate DNA damage response.

## Introduction

The DNA molecule is constantly exposed to endogenous and exogenous genotoxins, leaving it subjected to a multitude of lesions that, if left unchecked, can lead to genomic instability, a hallmark of cancer and other diseases (Jackson and Bartek 2009). Methyl methanesulfonate (MMS) is a widely used genotoxic agent that induces DNA damage by alkylating DNA bases, resulting in the formation of DNA adducts. The most prevalent adducts formed by MMS are N7-methylguanine (N7-MeG) and N3-methyladenine (N3-MeA) (Beranek 1990), while N1-methyladenine (N1-MeA) and N3-methylcytosine (N3-MeC) can also be generated during DNA replication (Wyatt and Pittman 2006). These DNA adducts can interfere with DNA synthesis and replication by inducing replication fork stalling and subsequent DNA strand breaks (Branzei and Foiani 2010). Additionally, MMS has been shown to cause damage to mitochondrial DNA, which can disrupt oxidative phosphorylation and contribute to increased generation of reactive oxygen species (ROS) (Salmon *et al*. 2004; Kitanovic and Wölfl 2006; Kitanovic *et al*. 2009) Under such circumstances, a comprehensive DNA Damage Response (DDR) is activated to tackle the consequences of DNA lesion accumulation, such as through DNA repair, telomere maintenance, chromatin remodeling, inhibition of DNA synthesis and cell cycle progression, transcription reprogramming, among others (Hanawalt 2015; Lanz *et al*. 2019; Cussiol *et al*. 2020).

Although DDR has been extensively studied at the molecular level, recent studies have proposed that the DDR is connected with metabolism of biomolecules such as carbohydrates and lipids (Simpson-Lavy *et al*. 2015; Yi *et al*. 2017; Ferrari *et al*. 2017). Remarkably, the existence of crosstalk between the DNA damage response (DDR) and phospholipid metabolism in eukaryotes was proposed several years ago (Zewail *et al*. 2003). Supporting this notion, numerous proteins involved in the phosphatidylinositol (PI) pathway were found to undergo phosphorylation in response to DNA damage induced by MMS (Zhou *et al*. 2016; Lanz *et al*. 2021). Inositol metabolites such as inositol polyphosphates (IPs) and inositol pyrophosphates (PP-IPs) have been implicated in cell cycle regulation and DNA damage repair (Banfic *et al*. 2013, 2016; Jadav *et al*. 2013), but the molecular basis for these effects is unknown. Furthermore, it was proposed that phosphatidylinositol phosphate lipids (PIPs) are enriched in the nucleus after DNA damage, serving as important mediators of ATR signaling in mammalian cells (Wang *et al*. 2017). Nonetheless, little is known about how DNA damage can regulate inositol metabolism and vice-versa.

In budding yeast, inositol biosynthesis is fine-tuned by the transcriptional repressor Opi1. In absence of inositol, Opi1 is localized to the perinuclear ER (endoplasmic reticulum) membrane establishing interactions with the integral membrane protein Scs2 and the phospholipid precursor phosphatidic acid (PA) (Craven and Petes 2001; Brickner and Walter 2004; Gaspar *et al*. 2017; Hofbauer *et al*. 2018). As a result, the heterodimeric transcriptional activator Ino2-Ino4 binds to a cis-acting inositol-sensitive upstream activation sequences (UAS_INO_) and upregulates expression of several genes related to phospholipid metabolism including the inositol-3-phosphate synthase (*INO1*), which promotes inositol *de novo* synthesis. Full repression of genes containing UAS_INO_ is also dependent on choline, leading to the designation of this sequence as inositol/choline responsive elements (ICRE) (Schüller *et al*. 1992). Once intracellular inositol concentration increases, PA is redirected to the synthesis of PI leading to its exhaustion. Subsequently, Opi1 migrates to the nucleus where it binds to the heterodimeric transcriptional factor Ino2-Ino4 and promotes transcriptional repression through the interaction with the proteins Sin3 and Cyc8, creating a scaffold for the recruitment of histone deacetylase (HDAC) complexes (Wagner *et al*. 2001; Jäschke *et al*. 2011; Kliewe *et al*. 2017) (Figure 1A). Importantly, deletion of Opi1 leads to constitutive expression of *INO1* with overproduction of inositol even when cells are supplemented with inositol (Graves and Henry 2000). Moreover, cells lacking Opi1 show constitutive activation of a large number of genes, most of which are regulated by Ino2-Ino4 (Santiago and Mamoun 2003; Jesch *et al*. 2005; Hoppen *et al*. 2005). Many of these genes are involved in phospholipid biosynthesis, although UAS_INO_ motifs are found in several genes related to other metabolic processes, which implicates Ino2-Ino4 in the control of the expression of distinct biological processes (Wimalarathna *et al*. 2011). Interestingly, phosphoproteomic analysis in budding yeast showed that the transcriptional repressor Opi1 is phosphorylated in a Mec1/Tel1-dependent manner after MMS exposure (Balint *et al*. 2015; Bastos de Oliveira *et al*. 2015; Lanz *et al*. 2021), which might suggest that Opi1 is involved in the DDR.

**Figure 1.**
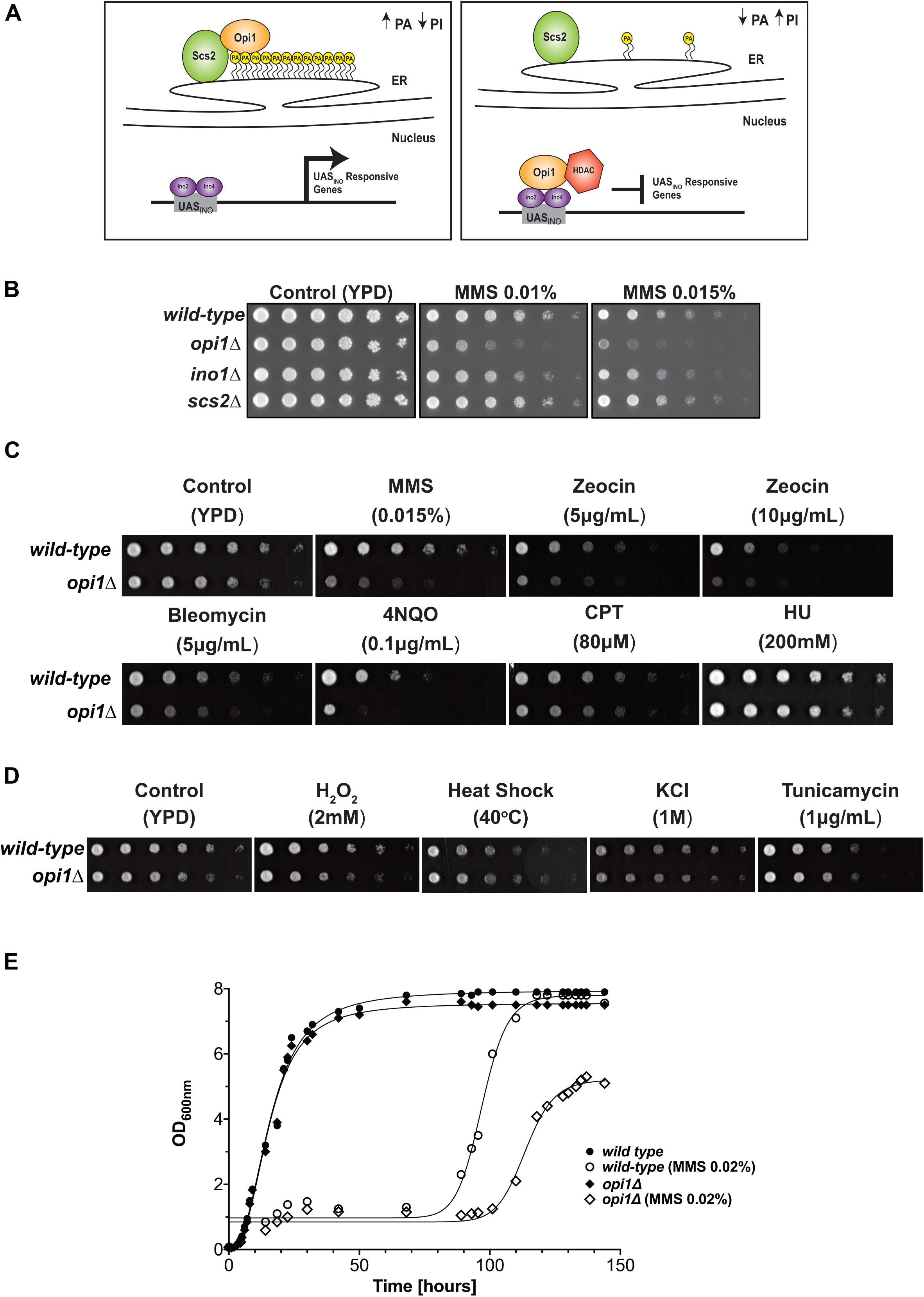
Cellular response of Opi1-deficient cells to genotoxic and proteotoxic stress. (**A**) Working model for how Opi1 repress transcription of target genes. (**B**) Effect of deletion of genes involved in inositol metabolism in sensitivity to genotoxins. (**C**) Cells lacking Opi1 show sensitivity to different genotoxins. (**D**) Cells lacking Opi1 does not show sensitivity to proteotoxic stress agents. (**E**) Growth rate of wild-type and *opi1Δ* cells, determined by monitoring optical density (OD_600_) over time. The data points were fitted to a nonlinear regression curve to analyze the growth kinetics. For (B, C and E), fourfold serial dilutions were spotted on the media indicated in the figure and plates were grown for 2–3 days at 30°C.

In this study, we employ an integrative approach combining RNA-seq transcriptome analysis, yeast genetics, molecular biology, cell biology, and biochemistry techniques to investigate the role of the transcriptional repressor Opi1 in the response to genotoxic stress. Our findings demonstrate that under genotoxic stress, cells lacking Opi1 exhibit enhanced sensitivity and cell cycle defects characterized by a delayed G1 to S-phase transition, resulting in reduced histone H2A phosphorylation. Notably, RNA-seq analysis reveal that during MMS-induced genotoxic stress, Opi1 is important to modulate expression of genes involved in important biological processes. Additionally, our study reveals that treatment with MMS leads to an increase in mitochondrial DNA (mtDNA) instability in *opi1Δ* cells. This observation suggests a potential link between mtDNA instability and the genotoxin sensitivity exhibited by cells lacking Opi1. Importantly, deletion of the transcriptional activator Ino2-Ino4 rescues the MMS sensitivity of *opi1Δ* cells showing that constitutive activation of Ino2-Ino4 is the cause of MMS sensitivity. Finally, while deletion of the inositol pyrophosphate kinase Kcs1 in an *opi1Δ* strain rescues MMS sensitivity and cell cycle defects, overexpression of Kcs1 in wild-type cells phenocopies the genotoxin sensitivity of an *opi1Δ* strain. These findings suggest that inositol pyrophosphates (PP-IPs) may counteract the function of Opi1 during MMS-induced genotoxic stress in yeast.

Collectively, these results highlight the crucial role of Opi1 as a key sensor of genotoxic stress and emphasize the significance of modulating PP-IPs synthesis as an integral component of the DNA damage response (DDR) and a critical factor for proper cell function in budding yeast.

## Material and Methods

### Yeast strains and plasmids

Strains generated in this study are isogenic from BY4741 and S288C (where indicated). Detailed information on all yeast strains and plasmids used in this study can be found in Table S1 and Table S2, respectively. To generate knockout and epitope-tagged strains, the one-step gene disruption method, as described by Rothstein (Rothstein 1983, 1991; Longtine *et al*. 1998), was employed. All yeast transformations were performed using the lithium acetate method (Gietz *et al*. 1992; Gietz and Woods 2006). PCR genotyping using specific primers (Table S3) was conducted to confirm the successful generation of knockout strains, while western blotting was performed to verify the presence of epitope tags in the respective strains. For plasmid construction, restriction digestion cloning was employed as follows: A PCR product containing 250bp of the *OPI1* promoter + the *OPI1* ope*n* reading frame (ORF) with s 3×HA sequence fused to the C-terminus and the *ADH1* terminator sequence was amplified from genomic DNA of a yeast strain expressing Opi1-HA and subsequently cloned into the pRS416 vector using XhoI and SacII restriction enzymes (New England Biolabs). To overexpress *INO1* and *KCS1*, the ORF encoding for *INO1* and *KCS1* were PCR amplified from yeast genomic DNA using specific primers. The amplified fragments were then cloned into a modified pYES2-NTC plasmid (a gift from Dr. Marcus Smolka), containing the *GAL1* promoter, using EcoRI and XhoI restriction sites (New England Biolabs).

### Growth conditions

Cells were grown in YPD (1% yeast extract, 2% peptone, 2% dextrose) and plasmid-bearing cells were grown in synthetic complete media lacking uracil (SC-URA) (0.17% YNB w/o aminoacids – Difco), 0.5% ammonium sulfate (MP Biomedicals) 0.07% CSM-URA dropout mix (Sunrise Science), 2% dextrose (Sigma). For experiments performed in the absence of inositol, SC-INO medium was prepared using an YNB without inositol (MP Biomedical). For expression of *INO1* and *KCS1* under the control of the *GAL1* promoter, cells were grown overnight in SC-URA medium with 2% dextrose and were plated in SC-URA plates in the presence of 2% dextrose (Sigma) or 2% galactose (Difco). For galactose induction in liquid cultures, the strains were initially grown in SC-URA medium supplemented with lactate (pH 5.5) as describe by the Haber lab (Haber and Leung 1996). Once the cell density reached an OD_600nm_ of 1.0, galactose was introduced into the culture to achieve a final concentration of 2%. To evaluate growth under respiratory conditions, cells were cultured in either YPGal (1% yeast extract, 2% peptone, 2% galactose) or YPG medium (1% yeast extract, 2% peptone, 3% glycerol). For the detailed concentrations and durations of specific chemical agents, including genotoxins and other drugs, please refer to the respective figure legends associated with the experimental data.

### Serial dilution assays

Overnight cultures were inoculated in the appropriate medium to an optical density (OD_600nm_) 0.1 and cells were grown until they reached the logarithmic growth phase and were subsequently normalized to an OD_600nm_ of 1.0. Fourfold serial dilutions were spotted on yeast plates and grown for 1–3 days at 30°C in the presence or absence of specific chemical agents, including genotoxins and other drugs. The specific concentrations of these agents used in the assay are provided in the corresponding figure and/or figure legends, serving as a clear reference for the experimental setup. Following the incubation period, the plates were manually inspected for growth changes and digitalized in an Uvitec Alliance 4.7 imaging system.

### Colony forming unit assay

To evaluate the effect of Opi1 on yeast cell viability following genotoxic stress, a colony-forming unit (CFU) assay was conducted. Exponentially growing yeast cells were treated with 0.1% MMS at 30°C for either 60 or 90 minutes. After treatment, 1 mL of cells was harvested by centrifugation, washed with fresh YPD, and resuspended in 1 mL of YPD medium. The samples were then normalized to an optical density at 600 nm (OD_600nm_) of 1.0 and diluted 1:2500 in YPD. Subsequently, 100 μL of each dilution was plated in triplicate on YPD agar plates. The plates were incubated at 30°C for 48 hours to allow colony formation. Following incubation, the number of colonies was counted, and the results were expressed as the percentage of viable cells compared to untreated cells. To determine statistical significance, multiple t-tests were employed to compare the mean viability values of the different strains, with a significance threshold set at α = 0.05.

### Western blotting

Whole-cell extracts (WCE) were prepared as follows: A 50 mg frozen cell pellet was lysed by bead beating at 4°C in lysis buffer containing 50 mM Tris-HCl, pH 7.5, 0.2% Tergitol, 150 mM NaCl, 5 mM EDTA, 1 mM phenylmethylsulphonyl fluoride (PMSF), Complete, EDTA-free Protease Inhibitor Cocktail (Roche), and PhosSTOP (Roche). Cell lysates were then harvested by centrifugation at 14,000 RPM for 15 minutes at 4°C, and the supernatant was collected. The protein samples were denatured by adding 3× SDS sample buffer (containing 188 mM Tris-HCl, pH 6.8; 3% SDS; 30% glycerol; 0.01% bromophenol blue; and 1× Bond-Breaker TCEP solution from Thermo Fisher Scientific) and heating at 95°C for 10 minutes. The denatured samples were resolved on SDS-PAGE gels for subsequent Western blot analysis. Proteins were transferred onto PVDF membranes (Amersham Hybond P – GE Healthcare). Phosphorylated histone H2A (gamma-H2A), total histone H2A, Rad53 and epitope tagged proteins were probed using specific antibodies: anti-gamma H2A (Ab17353-Abcam, 1:2500 dilution in TBST and 5% non-fat dry milk), anti-histone H2A (ab188312, Abcam, 1:500 dilution in TBST and 5% non-fat dry milk), anti-HA (12CA5; Roche, 1:10000 in TBST and 2% non-fat dry milk), anti-Rad53 (ab104232, Abcam, 1:3000 dilution in TBST and 5% non-fat dry milk). Anti-PGK1 was used as a housekeeping control (22C5D8, Abcam, 1:15000 dilution in TBST and 2% non-fat dry milk). After primary antibody, PVDF membranes were incubated in the presence of ECL HRP-linked secondary antibody (mouse: NA931-GE or rabbit: NA934-GE, 1:10000 dilution in TBST and 2% non-fat dry milk). Blots were developed using the Amershan ECL Prime detection reagent and imaged in a Uvitec Alliance 4.7.

### Cell cycle synchronization

Yeast cells were synchronized in the G1 phase as described before (Jablonowski *et al*. 2015). In brief, cells were initially grown in YPD medium at 30°C until they reached the logarithmic growth phase. Subsequently, α-factor (Zymo Research) treatment was applied at a concentration of 30 ng/ml (for *bar1Δ* background strains) for 2 hours. To release the cells from G1 arrest, they were centrifuged and resuspended in fresh medium. In order to assess the impact of MMS-induced genotoxic stress on S-phase progression, MMS was added at a concentration of 0.033% (equivalent to 3 mM) during the release step. To induce and intra-S-phase arrest of the cell cycle, cells were exposed to MMS at a concentration of 0.033% for 2 hours which is sufficient to activate the intra-S-phase checkpoint (Shirahige *et al*. 1998). For recovery of cells following MMS treatment, cells were exposed to MMS at a concentration of 0.033% for 2 hours, harvested, and subsequently resuspended in fresh YPD medium.

### Flow cytometry

Cell cycle analysis was performed as described before (Jablonowski *et al*. 2015). In brief, logarithmic growing yeast cells were collected, harvested, fixed in 1 ml of 70% ethanol (Sigma-Aldrich), and incubated overnight at 4°C. Cells were then centrifuged, and residual ethanol was dried in a speed-vac. After that, samples were solubilized in sodium citrate buffer (LabSynth) (50 mM, pH 7.2) and sonicated (three cycles of 3 sec, amplitude 30%) to release cell clumps. Samples were then incubated with 100 μg of RNAseA (Invitrogen) for 1 h at 37°C followed by incubation with 500 μg of Proteinase K (Invitrogen) for 1 h at 42 °C. Then, 1 μL of SYTOX Green (Thermo Fisher Scientific) was added to the samples and incubated for 2 h at 4°C protected from the light. Data were acquired using a BD Accuri C6 Flow Cytometer. For viability assessment, wild-type and *opi1Δ* cells were cultured in YPD liquid medium until the logarithmic phase of growth and treated with 0.1% MMS for 4 hours. After treatment, the samples were normalized to a concentration of 1×10^6^ cells/mL, harvested, and resuspended in 1 mL of 50 mM sodium citrate. Next, 0.4 μM propidium iodide (Thermo Fisher Scientific) was added, and the cells were immediately analyzed using a BD Accuri C6 Flow Cytometer. To distinguish between dead and live cells, we used boiled cells (100°C for 10 minutes) as a control for dead cells, and untreated cells to obtain a population of live cells. The data were obtained from three biological replicates and multiple t-tests were used to determine statistical significance, comparing the mean viability values of the strains with the significance set at α = 0.05.

### Real Time Quantitative PCR (RT-qPCR) analysis

Cells were grown overnight in YPD and diluted next day to an OD_600nm_ of 0.1 in synthetic complete medium either supplemented or lacking inositol (SC+INO or SC-INO). Cells were grown for at least two cell cycle divisions and immediately harvested at 4°C, washed with sterile ultra-pure H_2_O, and kept at –80°C. Alternatively, log phase cells were treated with MMS (Sigma-Aldrich) 0.1% for 1 hour. Total RNA from 5×10^7^ cells was extracted using RNeasy mini kit (Qiagen). One μg total RNA was used for cDNA synthesis using Quantinova Reverse Transcription kit (Qiagen). Real time PCR was performed in an ABI Prism 7500 (Applied Biosystems) using Quantinova Probe PCR kit (Qiagen) in the presence of Taqman^®^ probes for *INO1* (Sc04136910_s1) and *ACT1* (Sc04120488_s1) (Thermo Fisher Scientific). The *ACT1* gene served as an internal standard for normalization as it was previously shown that there is no change in mRNA levels either after inositol depletion (Ye *et al*. 2013; Gaspar *et al*. 2017) or MMS treatment (Gasch *et al*. 2001). Relative values of mRNA are described as fold change relative to a control condition indicated in the experiment. The data were obtained from three biological replicates and statistical analysis was performed using a one-way analysis of variance (ANOVA) with Tukey’s post hoc test to determine the significant differences between the groups.

### RNA-Seq sample preparation and analysis

Cells were grown overnight in SC + INO media and diluted next day to an OD_600nm_ of 0.1 in 250 mL of SC + INO media in biological duplicates. Cells were grown for at least two cell cycle divisions (∼ OD_600nm_ = 0.4) then vacuum filtered in 0.22 µm nitrocellulose membranes to remove media and immediately “flash frozen” in liquid nitrogen. 1 mL of frozen 1X RIPA buffer (Millipore) was combined into each frozen pellet and lysed by mechanical grinding with a Spex 6750 Freezer Mill at a rate of 10 Hz for 2 minutes. The lysate was then thawed on ice and centrifuged at 2,000 g for 2 minutes at 4°C to remove debris. 250 µL of cleared lysate was used for RNA extraction by combining it with 250 µL of 2X AE Buffer (100 mM sodium acetate pH 5.2, 20 mM EDTA), 50 µL of 10% SDS and 500 µL of acid phenol: chloroform pH 5.2. The mixture was incubated at 65°C for 10 minutes under 1400 rpm agitation, then transferred to ice for 5 minutes. The solution was then spun at 15,000 g for 15 minutes at 4°C. The supernatant was then transferred to new microcentrifuge tubes and 550 µL of chloroform was added, followed by centrifugation at 15,000 g for 5 minutes at 4°C. The aqueous phase was then transferred to new microcentrifuge tubes and RNA was precipitated in 0.3M sodium acetate and 400 µL of isopropanol after centrifugation at 15,000 g for 20 minutes at 4°C. RNA pellets were washed with 75% ethanol and centrifuged at 15,000 g for 5 minutes at 4°C. Finally, RNA pellets were resuspended in nuclease-free water and submitted to Bioanalyzer High Sensitivity DNA Assay (Agilent) for quality check and quantified using Qubit RNA High Sensitivity (Invitrogen). 1 µg of total RNA was used as input for preparing the libraries for RNA sequencing with Ultra RNA Library Prep II (New England Biolabs), following the recommended workflow. Final cDNA libraries after PCR were quantified by Qubit dsDNA High Sensitivity (Invitrogen). Samples were then equimolarly pooled and distributed on Illumina NextSeq High 500/550, resulting in ∼ 7-8 million reads per sample. Reads were demultiplexed using bcl2fastq (version 2.20) and quality of reads was assessed using FastQC (version 0.11.8). Reads were then aligned to the S288C reference genome (release R64-2-1) using STAR (version 2.5.4b) (Dobin *et al*. 2013). Differential gene expression was calculated with DESeq2 (version 1.40.0) (Love *et al*. 2014) using the design “∼ genotype + condition + genotype: condition”. A threshold of 0.8 log_2_ fold change and adjusted p-value of 0.05 was used to analyze differentially expressed genes between samples. A principal component analysis (PCA) plot for the two biological replicates across genotypes and treatments is shown in Supplementary Figure 1. Table S4 (“genotype_OPI1_vs_WT.csv”) shows the differentially expressed genes between the *opi1Δ* and wild-type strains in untreated condition. Table S5 (“treatment_MMS_vs_unt.csv”) shows differentially expressed genes in the wild-type strain treated with MMS relative to untreated. Table S6 (“interaction.csv”) shows the differential effect of MMS treatment in *opi1Δ* strain relative to the wild-type (interaction term). Table S7 (“treatment_MMS_vs_unt_OPI1.csv”) shows the differentially expressed genes in the *opi1Δ* strain treated with MMS relative to the *opi1Δ* without treatment. Table S8 (“OPI1_vs_WT_treatment.csv”) shows the differentially expressed genes between the *opi1Δ* and the wild-type strains both treated with MMS.

To identify enriched biological processes and cellular components among the list of differentially expressed genes (DEGs), we utilized the PANTHER database (http://pantherdb.org) and the Saccharomyces Genome Database (SGD) with the following parameters: PANTHER version 17.0 Overrepresentation Test, FISHER test with FDR correction (Thomas *et al*. 2022), and the Gene Ontology Slim term mapper (https://www.yeastgenome.org/goSlimMapper), respectively. Additionally, interaction networks of differentially expressed gens (DEGs) were generated using the STRING v11.5 database (http://string-db.org) (Szklarczyk *et al*. 2019).

### Respiratory capacity assay

Oxygen consumption rates were measured using a modified protocol based on the method described by (Zulkifli *et al*. 2020) using a high-resolution O2K Fluorespirometer (Oroboros, Innsbruck, Austria). For each assay, 10^6^ cells were added to the Oroboros chamber containing a final volume of 2.0 ml of YPEG medium at 30°C under agitation. Briefly, exponentially growing cells in YPD were harvested, resuspended in respiratory medium (YPEG; 2% glycerol and 1% ethanol), and incubated for at least 3 hours at 30°C. Finally, cells were transferred to O2K chambers for measurement of O_2_ consumption rates. After basal respiration assessment, 5μM of the mitochondrial uncoupler FCCP (Sigma-Aldrich) was added to measure maximal respiration. At the end, 2μM of antimycin A (Sigma-Aldrich) was added to inhibit mitochondrial respiration. Non-mitochondrial oxygen consumption rates were subtracted from all measurements. Optimal concentrations of FCCP and antimycin A were previously determined by titration. The data were obtained from two biological replicates and statistical analysis was performed using a one-way analysis of variance (ANOVA) with Tukey’s post hoc test to determine the significant differences between the groups.

### Microscopy analysis

Images were acquired in a Zeiss LSM 780 microscope. Laser lines at 561nm were used for the detection of Opi1-GFP tagged protein in yeast cells. For yeast Opi1-GFP analysis, cells were grown in synthetic complete media (SC +INO) until the log phase (OD = 0.3) and shifted to SC –INO for 1 hour. Finally, MMS (0.1%) was added to the cells and incubated for 30 min at 30°C. At each condition, cells were washed in sterile water and resuspended in fresh synthetic complete media lacking tryptophan (SC-TRP). Live yeast cultures were mounted on an agarose slide pad (1.2% agarose in SC-TRP media) and >100 cells were scored for each replicate. Cells were manually counted for Opi1-GFP localization (nuclear, transition from the nucleus to the ER and ER localization). The data were obtained from two biological replicates and statistical analysis was performed using a one-way analysis of variance (ANOVA) with Tukey’s post hoc test to determine the significant differences between the groups.

## Data availability

Strains and plasmids are available upon request. Tables S1, S2 and S3 contain the list of yeast strains, plasmids and primers used in this study, respectively. Table S4 to S8 contain the RNA-seq dataset as described in Material and Methods section. Figure S1 contains the Principal Component Analysis showing the variation among biological replicates in RNA-seq. Figure S2 contains supplemental data in support of Figure 1. Figure S3 contains supplemental data in support of Figure 2. Figure S4 contains supplemental data in support of Figure 3A and B. Figure S5 contains supplemental data in support of Figure 4. Figure S6 and S7 contains supplemental data in support of Figure 5. Figure S8 contains supplemental data in support of Figure 5D and E. Figure S9 contains supplemental data in support of Figure 6B.

**Figure 2.**
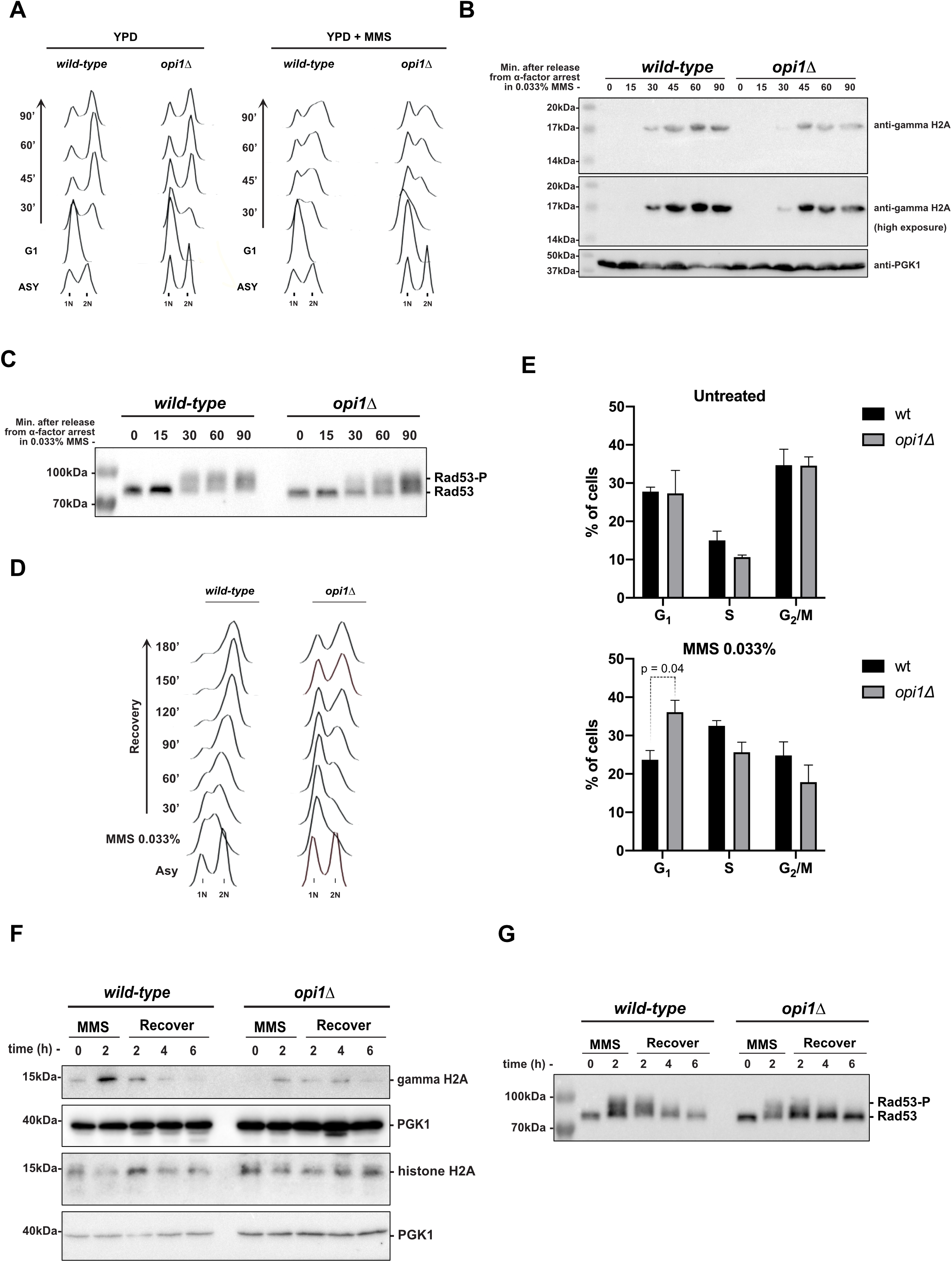
Cells lacking Opi1 show a delayed G1 to S-phase transition and activation of the DDR upon genotoxic stress induced by MMS. (**A**) Flow cytometry analysis depicting S-phase progression in wild-type and *opi1Δ* strains, (**B**) Western-blot analysis of gamma-H2A (histone H2A phosphorylation) in the indicated strains. (**C**) Western-blot analysis showing MMS-induced Rad53 phospho-shift in the indicated strains. (**D**) Flow cytometry analysis illustrating the resumption of S-phase progression in wild-type and *opi1Δ* strains after MMS treatment (0.033% for 2 hours). (**E**) Quantification of the percentage of cells in G1, S, or G2/M phases in untreated cells and after MMS treatment (0.033% for 2 hours). The results shown are representative of three independent experiments, and error bars indicate the standard deviation. Two-way ANOVA with Tukey’s post-test was used for significance analysis between samples. (**F**) Western-blot analysis of gamma-H2A and total histone H2A in the indicated strains. (**G**) Western-blot analysis showing MMS-induced Rad53 phospho-shift in the indicated strains. Note: For A-C, cells were arrested in G1 after 2 hours treatment with α-factor and then released in fresh YPD medium ± 0.033% MMS. All strains are *bar1Δ*. For D-G, cells were treated with 0.033% MMS for 2 hours and then release in fresh YPD medium. western-blots were probed with anti-gamma-H2A, anti-histone H2A, anti-Rad53 and anti-Pgk1 (loading control) antibodies as described in *Material and methods*.

**Figure 3.**
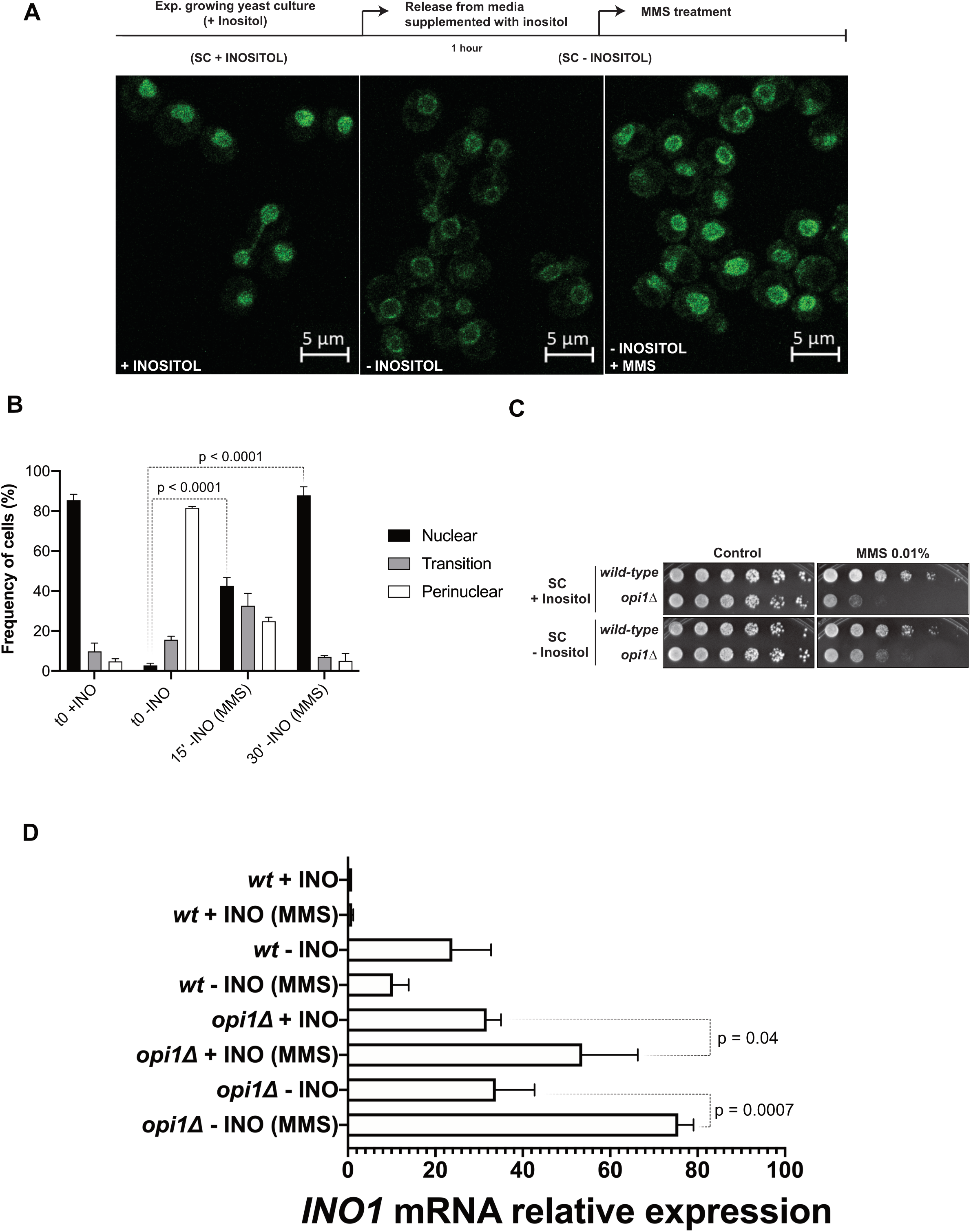
Opi1 translocates to the nucleus after MMS-treatment and is required to repress *INO1* expression during genotoxic stress. **(A-B)** Effect of inositol availability and MMS treatment on Opi1-GFP localization. Cellular localization of Opi1-GFP was scored manually and the means and standard deviations from two independent experiments are shown. (**C**) Cells treated with MMS in medium lacking inositol are still viable. (**D**) Effect of inositol availability and MMS treatment on *INO1* expression by RT-qPCR. Relative values of mRNA are described as fold change relative to a control condition (*wt* + INO). For C, fourfold serial dilutions were spotted on the media indicated in the figure and plates were grown for 2–3 days at 30°C. For D, experiments were made with three biological replicates and statistical significance of the differences observed was calculated using one-way ANOVA with Tukey’s multiple comparisons test.

**Figure 4.**
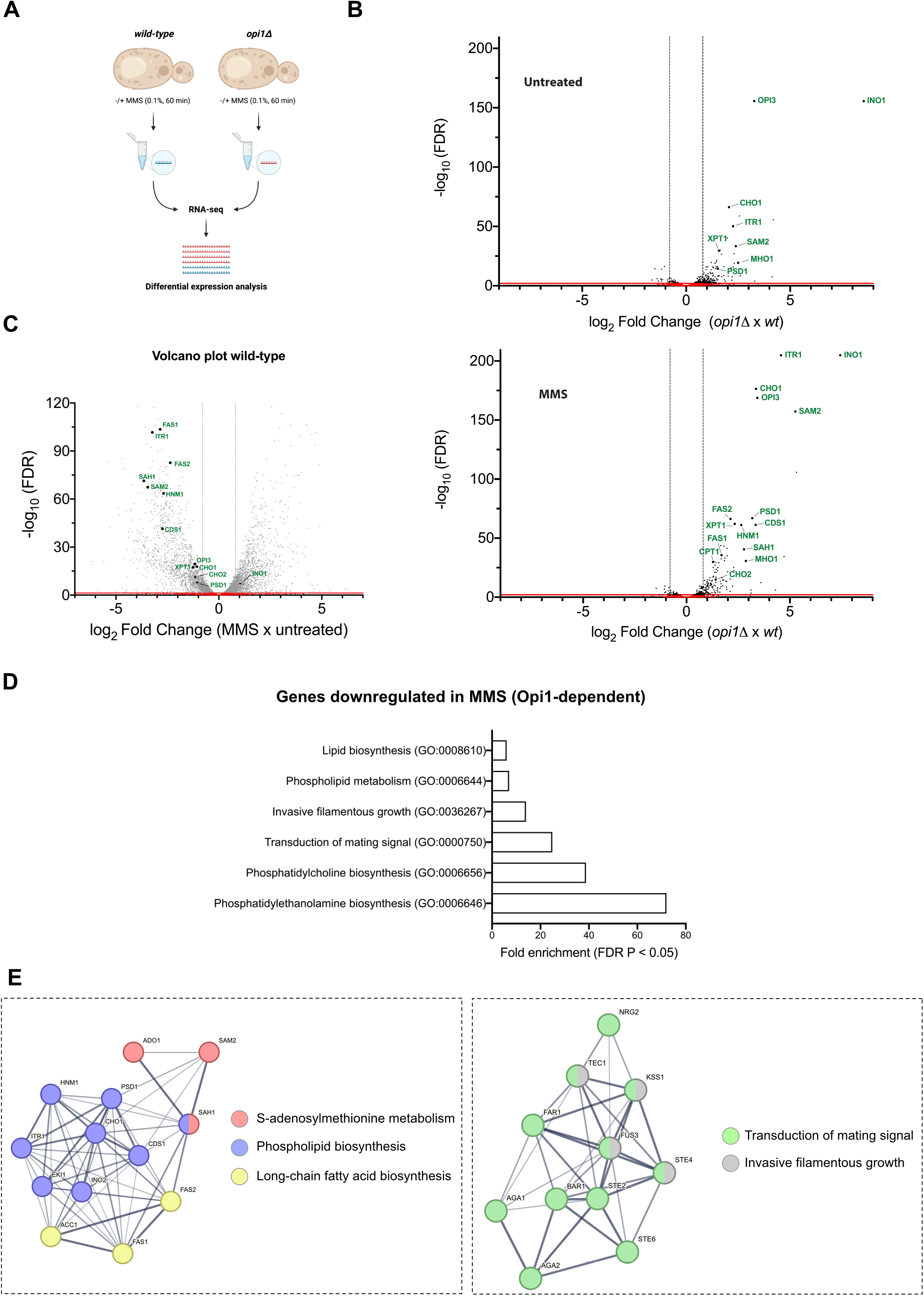
Opi1 is important to modulate gene expression during MMS-induced genotoxic stress. (**A**) Schematic representation of RNA-seq performed in wild-type and opi1Δ cells cultured in minimal medium (SC + INO) with or without 0.1% MMS treatment for 60 minutes. Created with BioRender.com. (**B**) Volcano plot illustrating the differential gene expression between *opi1Δ* vs wild-type cells (top panel) and between *opi1Δ* + MMS vs wild-type + MMS cells (bottom panel). (**C**) Volcano plot displaying the differential gene expression between wild-type + MMS vs wild-type cells. For B and C, genes with significant differential expression are plotted based on a threshold of ±0.8 log_2_ fold change and an adjusted p-value of 0.05. The red line (x-axis) and dotted line (y-axis) represent the significant cut-off. Known genes regulated by Opi1 are labeled in both plots. (**D**) Bar plot illustrating the top enriched biological processes identified in the gene set that are repressed during MMS treatment in an Opi1-dependent manner (Table S6). Each bar represents a specific biological process, and the height of the bar corresponds to the fold enrichment with a p-adjusted value of 0.05. Gene ontology (GO) analysis was conducted in PANTHER using the overrepresentation test with *Saccharomyces cerevisiae* serving as the reference list, which includes all genes in database. The test type used was FISHER, and the false discovery rate (FDR) correction was applied. (**E**) Gene Ontology (GO) term enrichment network depicting the relationships between genes grouped in clusters that are repressed during MMS treatment in an Opi1-dependent manner (Table S6). The enrichment analysis was performed using the STRING database, which integrates various sources of functional information with a confidence score >0.4 (medium confidence). Nodes in the network represent genes, while edges indicate the connections between them. The color of the nodes represents a biological process, and the thickness of the edges represents the increased confidence of the interaction between nodes.

**Figure 5.**
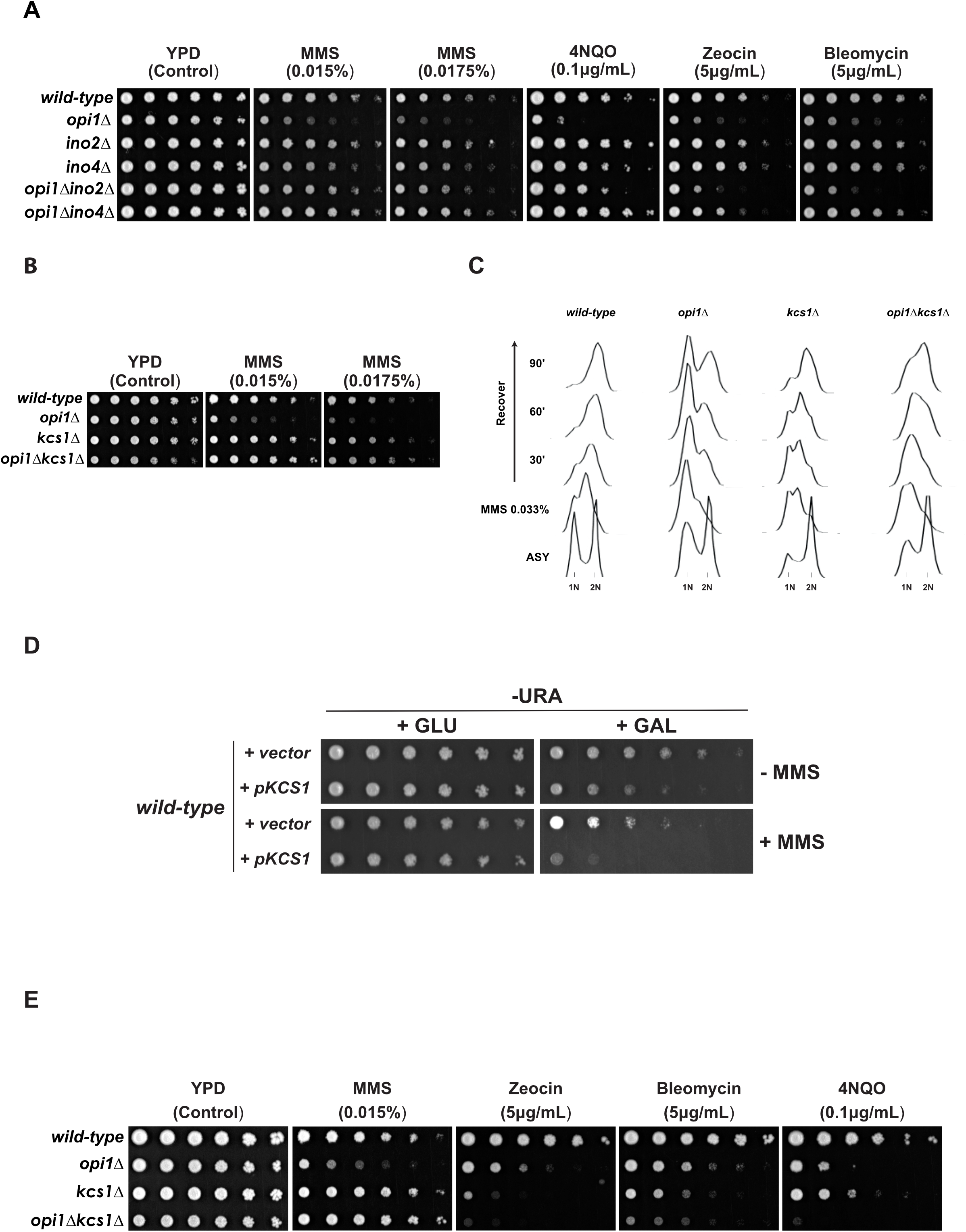
Opi1 counteracts the function of the transcription repressor Ino2-Ino4 and the inositol pyrophosphate kinase Kcs1 during MMS-induced genotoxic stress. (**A**) Deletion of *INO*2 and *INO*4 rescues the genotoxin sensitivity of cells lacking Opi1. **(B-C)** Deletion of the inositol pyrophosphate kinase *KCS1* rescues the MMS sensitivity (B) and the delayed G1 to S-phase transition (C) of cells lacking Opi1. (**D**) Overexpression of Kcs1 phenocopies the MMS sensitivity of an *opi1Δ* strain. (**E**) Different from the effect on MMS, deletion of Kcs1 causes a synergistic increase in sensitivity of *opi1Δ* cells in the presence of several DNA damage agents. For A, B, D and E fourfold serial dilutions were spotted on the media indicated in the figures and plates were grown for 2–3 days at 30°C.

**Figure 6.**
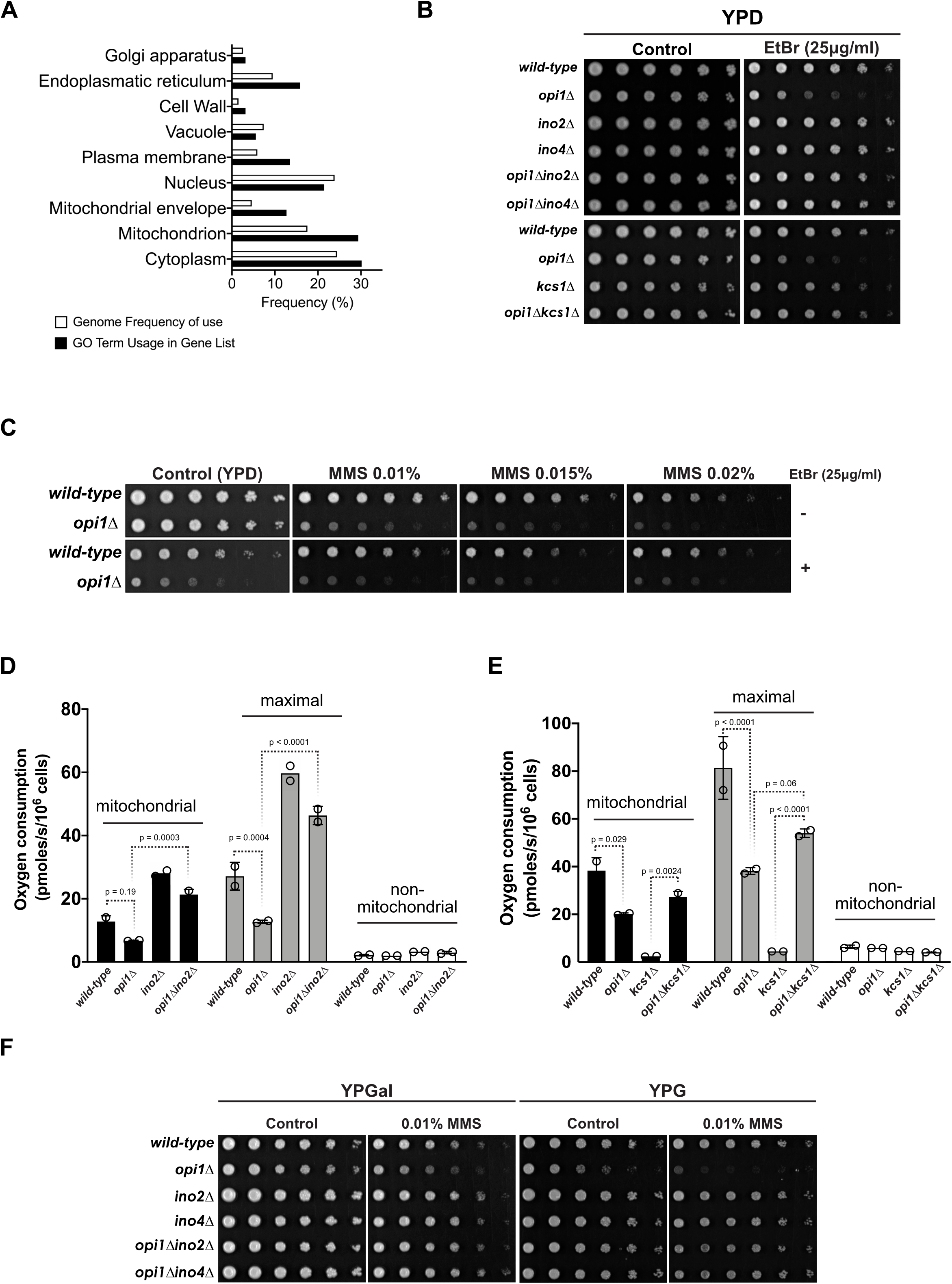
Opi1 is important to prevent mitochondrial DNA instability and to promote respiratory function during MMS-induced genotoxic stress. (**A**) Gene Ontology (GO) Slim term analysis using the Yeast Genome Database’s GO Slim Mapper revealing enrichment of genes associated with specific cellular components that are upregulated in cells lacking Opi1 upon MMS treatment (Table S8). All genes from *Saccharomyces cerevisiae* were included as a reference list. (**B**) Deletion of *INO*2, *INO*4 and *KCS1* rescues the ethidium bromide (EtBr) sensitivity of cells lacking Opi1. (**C**) Combined treatment of MMS and EtBr does not synergistically enhance the sensitivity phenotype of an *opi1Δ* strain beyond the sensitivity conferred by each drug independently. (**D**) Deletion of *INO*2 completely restores mitochondrial respiratory capacity of cells lacking Opi1. (**E**) The *opi1Δkcs1Δ* double mutant exhibits restored mitochondrial respiratory capacity, which is impaired in the single *opi1Δ* and *kcs1Δ* mutants. (**F**) Deletion of *INO*2, *INO*4 rescues the MMS sensitivity of cells lacking Opi1 in medium containing respiratory carbon sources (galactose and glycerol). For B, C, and E, fourfold serial dilutions of the samples were spotted onto the designated growth medium as indicated in the figure. The plates were then incubated at 30°C for 2-3 days, except for plates containing YPG medium, which were photographed after 4 days of incubation.

## Results

### Opi1 is important for cell survival during genotoxic stress

To determine whether cells lacking Opi1 are more sensitive to DNA damage, we performed sensitivity assays in the presence of genotoxins. We found that *opi1Δ* cells showed sensitivity to genotoxic stress induced by the DNA-alkylating agent MMS (Figure 1B), indicating that Opi1 is important for the response to genotoxic stress. To confirm that this sensitivity to MMS was specific to *OPI1* deletion, we constructed a pRS416 plasmid expressing Opi1-HA from its endogenous promoter and show that Opi1-HA expression rescues the MMS sensitivity in *opi1Δ* (Supplementary Figure 2A and B respectively). Furthermore, deletion of genes that counteract Opi1 repressing functions, such as *INO1* and *SCS2*, did not lead to MMS sensitivity (Figure 1B).

Besides MMS, we performed a screening of several known DNA-damaging agents to assess for genotoxicity in *opi1Δ* cells (Figure 1C). Interestingly, we found that *opi1Δ* cells are sensitive to different genotoxins such as the radiomimetic drugs bleomycin and zeocin and the UV-mimetic 4NQO, while they did not show sensitivity to the replication stress inducers hydroxyurea (HU) and camptothecin (CPT). Importantly, cells lacking Opi1 did not shown sensitivity to agents that activate the ESR (<u>E</u>nvironmental <u>S</u>tress <u>R</u>esponse) and the UPR (<u>U</u>nfolded <u>P</u>rotein <u>R</u>esponse) indicating that Opi1 shows specificity to cope with increased DNA damage inflicted by genotoxins (Figure 1D).

To determine if the increased sensitivity to genotoxins observed in cells lacking Opi1 is due to loss of viability rather than a decreased growth rate, we used a colony-forming unit assay and flow cytometry to monitor cell survival (Supplementary Figure 2C and D respectively). Our results show that *opi1Δ* cells do not exhibit a significant decrease in cell viability compared to wild-type cells, indicating that the hypersensitivity to genotoxins is likely due to delayed cell proliferation rather than cell death. Consistent with that, *opi1Δ* cells show growth defect in YPD liquid medium in the presence of MMS 0.02% suggesting an increased arrest of the cell cycle in *opi1Δ* cells (Figure 1E).

### Cells lacking Opi1 show delayed G1 to S-phase progression and decreased levels of gamma-H2A

Since *opi1*Δ mutants are sensitive to genotoxic stress (Figure 1C) and show a prolonged arrest of growth (Figure 1E), we sought to investigate if these cells have defects in the DDR. To investigate the potential involvement of Opi1 in the regulation of the DNA damage response (DDR), we subjected asynchronous yeast cells to genotoxic agents, such as MMS, Zeocin, and 4NQO, and monitored the activation of Rad53, a pivotal kinase in the signal transduction pathway responsible for DDR regulation (Pellicioli and Foiani 2005; Branzei and Foiani 2006; Cussiol *et al*. 2020). Our findings revealed no significant differences in Rad53 activation between wild-type and *opi1Δ* cells in asynchronous conditions, suggesting that Opi1 may not play a critical role in the regulation of the Rad53 axis within the DDR (Supplementary Figure 3).

Given the indications from our results that genotoxic stress leads to delayed cell proliferation in *opi1Δ* cells (Figure 1E), we proceeded to investigate the role of Opi1 in the regulation of the cell cycle. To accomplish this, we utilized flow cytometry to monitor cell cycle progression. Initially, we arrested cells in G1 phase using α-factor and subsequently released them into fresh YPD medium in the presence or absence of MMS 0.033%, a concentration known to reduce S-phase progression and induce a robust activation of the intra-S-phase checkpoint (Tercero and Diffley 2001). Our findings reveal that cells lacking Opi1 exhibit a noticeable delay in S-phase progression under both unchallenged conditions and in the presence of MMS, as depicted in Figure 2A. Furthermore, it is noteworthy that cells lacking Opi1 consistently display a discernible G1 peak, indicating a G1/S arrest in the cell cycle after release from pheromone arrest. Interestingly, cells lacking Opi1 show a delayed phosphorylation of histone H2A (gamma-H2A) and a reduced intensity of the gamma-H2A signal at later time points (Figure 2B). Following DNA damage, the sensor kinases Mec1 and Tel1 catalyze the phosphorylation of histone H2A at serine residue 129 (gamma-H2A) in extensive regions surrounding the DNA lesion, creating a platform for the recruitment of various DNA damage response proteins (Rogakou *et al*. 1998; Downs *et al*. 2000), thus serving as a readout for DDR activation (Fernandez-Capetillo *et al*. 2004). Importantly, a previous study has demonstrated that gamma-H2A is enriched in regions containing newly replicated DNA, particularly behind the replication forks at early-firing replication origins following MMS treatment (Balint *et al*. 2015). The observed decrease in gamma-H2A levels in cells lacking Opi1 may be attributed to a decrease in the firing of replication origins. Importantly, our results also reveal a delayed activation of Rad53 in Opi1-deficient cells (Figure 2C), which aligns with the experimental observations regarding gamma-H2A (Figure 2B). Specifically, we observed the appearance of a phosphorylated isoform of Rad53 at 30 minutes of MMS treatment in wild-type cells, while this band was not detected at this time point in *opi1Δ* cells (Figure 2C).

Next, we assessed the ability of cells to recover from genotoxic stress induced by MMS. Exponentially growing cells were treated with 0.033% MMS for 2 hours to induce an intra S-phase arrest (Gasch *et al*. 2001). Afterward, cells were released into fresh medium, and cell cycle progression and DDR deactivation were monitored. We observed that MMS-treated *opi1Δ* cells had a significant increased population of G1 cells, indicating a persistent G1/S arrest of the cell cycle (Figure 2D and E). In contrast, wild-type cells displayed an intra-S phase arrest, which is consistent with the replication stress induced by MMS (Figure 2D). It is noteworthy that, consistent with the result from Figure 2B, gamma-H2A levels were significantly lower in *opi1Δ* cells relatively to wild-type after 2 hours of MMS treatment. However, no significant difference was observed in the kinetics of gamma-H2A disappearance during the recovery period (Figure 2F). Notably, the decrease in gamma-H2A levels observed in *opi1Δ* cells was not due to histone depletion since total histone H2A levels were not decreased in *opi1Δ* cells after genotoxic stress (Figure 2F). Also, we found that both wild-type and *opi1Δ* cells exhibited comparable levels of Rad53 activation after 2 hours of MMS treatment, and there was no difference in kinetics of Rad53 deactivation during recovery (Figure 2G). Our results suggest that DNA damage signaling downregulation is not altered in *opi1Δ* cells but indicate that MMS treatment induces cell cycle defects related to a delayed G1 to S-phase progression.

### Opi1 migrates to the nucleus upon genotoxic stress induced by MMS

Knowing that transcriptional repression function of Opi1 is dependent on its cellular localization (Figure 1A) and because its absence causes genotoxin sensitivity and cell cycle defects (Figure 1C and Figure 2 respectively), we investigated Opi1 localization after treatment with 0.1% MMS since this concentration was previously used to monitor global changes in gene expression (Jelinsky and Samson 1999). Using a strain expressing Opi1-GFP, we recapitulated Opi1 cellular localization during inositol metabolism (Figure 3A panels I and ii). MMS treatment of cells supplemented with 100μM of inositol (+INO), a concentration known to sufficient repress *INO1* (Hirsch and Henry 1986), did not cause any significant alteration in Opi1-GFP intensity or localization (data not shown). Surprisingly, MMS treatment of cells kept without inositol supplementation (– INO) induced a rapid translocation of Opi1-GFP to the nucleus (Figure 3A-B and Supplementary Figure 4). Since Opi1 mediates gene repression when it translocates to the nucleus, this observation might suggest that the nuclear localization of Opi1 is important to mediate with genotoxic stress resistance. Importantly, we found that yeast cells treated with MMS in SC-INO remained viable (Figure 3C), which implies that Opi1 does not fully repress *INO1* in these conditions, otherwise cells would die.

Subsequently, to understand if Opi1 migration to the nucleus in the presence of MMS causes gene repression, we looked at *INO1* expression during MMS treatment by RT-qPCR analysis (Figure 3D). While MMS treatment does not change *INO1* expression in wild-type cells supplemented with inositol (+INO), there is a substantial decrease in *INO1* expression in the absence of inositol (-INO). Strikingly, *INO1* expression significantly increases in an *opi1Δ* strain upon MMS treatment both in +INO and –INO conditions (Figure 3D). These results show that under genotoxic stress Opi1 migrates to the nucleus to repress *INO1* expression in order to downregulate inositol synthesis. Because Opi1 represses genes containing UAS_INO_, it is possible that in its absence, several other genes are also upregulated upon MMS treatment.

### Opi1 modulates gene expression during genotoxic stress induced by MMS

In order to gain a deeper understanding of the impact of Opi1-mediated gene expression regulation in response to MMS treatment on a transcriptome-wide scale, we conducted RNA-seq experiments using exponentially growing cells (wild-type vs *opi1Δ*) supplemented with 100μM inositol (SC + INO) in the presence of ± 0.1% MMS for 1 hour (Figure 4A). We selected this specific experimental condition based on two primary reasons: Firstly, our experiments have demonstrated the heightened sensitivity of Opi1-deficient cells to MMS when grown in medium supplemented with inositol (Figure 3C). Secondly, our own observation revealed that MMS-induced upregulation of *INO1* expression in *opi1Δ* cells is independent of inositol supplementation (Figure 3D).

Previous studies utilizing transcriptome analysis have characterized genes whose expression is modulated by Opi1 (Santiago and Mamoun 2003; Jesch *et al*. 2005). As anticipated, the results of our differential expression analysis revealed the upregulation of several canonical Opi1 targets in *opi1Δ* cells (Figure 4B top panel and Table S4). Interestingly, our results demonstrated that the expression of several Opi1 canonical targets was further augmented following MMS treatment in *opi1Δ* cells (Table S8 and Figure 4B, compare top and bottom panels), indicating the crucial role of Opi1 in modulating gene expression during such conditions. Moreover, to reinforce this hypothesis, we observed that genes highlighted in Figure 4B were repressed in wild-type cells exposed to MMS (Figure 4C and Table S5). This finding further supports the notion that Opi1 plays a critical role in regulating gene expression in response to MMS treatment and that its absence leads to misregulation of several genes, ultimately resulting in MMS sensitivity. Curiously, contrary to the observed trend in *INO1* expression determined by RT-qPCR (Figure 3D), *INO1* expression did not follow the same pattern as other Opi1 targets, as MMS treatment did not increase *INO1* expression in *opi1Δ* cells (compare Figure 4B, top and bottom panels). In fact, our data showed a 2-fold increase in *INO1* expression following MMS treatment (Figure 4C and Table S5). The observed discrepancy in *INO1* expression between the RT-qPCR and RNA-seq analyses could stem from various factors, including technical variability inherent to the methods, biological variability between samples, and differences in sensitivity and dynamic range. Further investigations and validation experiments are needed to understand the underlying reasons for this discrepancy in gene expression quantification.

To clarify the role of Opi1 during MMS treatment, we sought to identify genes exhibiting differential expression in an Opi1-dependent manner (Table S6). We then selectively screened genes showing a ± 0.8 log_2_ fold change with an adjusted p-value of ≤ 0.05 and conducted a statistical overrepresentation analysis using PANTHER database (http://pantherdb.org) to investigate biological processes that were enriched within our gene list. With that, we were able to see which biological process are been misregulated upon MMS treatment in an Opi1 dependent manner. Notably, we observe a significant enrichment of phospholipid biosynthesis genes within our dataset (Figure 4D), thereby corroborating the established role of Opi1 in downregulating phospholipid biosynthesis in yeast. Moreover, we observe that Opi1 is pivotal in repressing the expression of proteins associated with MAPK signaling such as cellular conjugation, transduction of mating signal (pheromone response), and filamentous growth (Figure 4D), thereby indicating its significance in modulating these biological processes during genotoxic stress. In addition, we generated an interactome of the differentially expressed genes which aligns in clusters using the STRING database (http://stringdb.org) (Figure 4E). Importantly, the upregulation of genes involved in the mating signaling pathway is associated with the inhibition of the G1 to S-phase transition. This finding is particularly relevant since we observed that cells lacking Opi1 exhibit a delayed transition from G1 to S-phase under genotoxic stress induced by MMS (Figure 2A, D, and E), which can be correlated with the observed delay in cell growth in both solid and liquid cultures (Figure 1C and E, respectively).

Finally, our findings challenge the canonical role of Opi1 as a gene repressor by also demonstrating its involvement in the upregulation of several genes related to amino acid biosynthesis, including arginine and histidine biosynthesis, the mitochondrial glycine cleavage system and sulfate assimilation (Supplementary Figure 5A and B). However, it is important to emphasize that we cannot exclude the possibility that Opi1 indirectly regulates many of these targets by influencing the expression of genes involved in transcriptional regulation.

### The constitutive activation of Ino2-Ino4 responsive genes is the cause of genotoxin sensitivity in cells lacking Opi1

A previous study by (Heyken *et al*. 2005) provided evidence for a physical interaction between Opi1 and a functional domain within the transcriptional factor Ino2. This interaction was shown to result in the repression of Ino2’s ability to promote gene expression. Moreover, our RNA-seq analysis revealed that *INO2* and *INO4* expression is upregulated in *opi1Δ* cells following MMS treatment (Table S8), indicating a role for Opi1 in *INO2* repression during genotoxic stress. In line with that, we investigated if *INO2* and *INO4* deletions could rescue the genotoxin sensitivity of cells lacking Opi1. Remarkably, *opi1Δino2Δ* and *opi1Δino4Δ* strains are rescued for the sensitivity to MMS and 4NQO, which indicates that constitutive expression of Ino2-Ino4 responsive genes is causing the hypersensitivity to these genotoxins (Figure 5A). Interestingly, our findings demonstrate that the deletion of *INO4* alleviates the sensitivity to zeocin and bleomycin, while the deletion of *INO2* does not have the same effect. Moreover, we notice that *ino4Δ* cells display sensitivity to heat shock, but not to other types of proteotoxic stress inducers (Supplementary Figure 6). Intriguingly, *ino2Δ* cells do not exhibit sensitivity to heat shock and the deletion of Opi1 rescues the heat shock sensitivity of *ino4Δ* cells while conferring sensitivity to *ino2Δ* cells (Supplementary Figure 6). These observations suggest that Ino2 and Ino4 can play independent roles in the transcriptional regulation during different types of stress. Importantly, our findings are supported by a previous study that examined gene expression under carbon and nitrogen limitation conditions, revealing distinct modulation patterns by Ino2 and Ino4, which target different sets of genes (Chumnanpuen *et al*. 2013). These consistent findings underscore the unique and context-dependent roles of Ino2 and Ino4 in transcriptional regulation during stress responses.

The transcriptional activator Ino2-Ino4 upregulate the expression of several other genes counteracting Opi1 suggesting that constitutive expression of another Ino2-Ino4 target in *opi1Δ* cells is causing MMS sensitivity. Thus, we conducted a screening of a small set of genes which expression is dependent on Ino2-Ino4 and that are upregulated in *opi1Δ* cells treated with MMS (Supplementary Table S8). We selected genes involved in phosphatidylinositol (*INO1* – inositol-3-phosphate synthase; *ITR1* – inositol transporter) and phosphatidylcholine biosynthesis (*OPI3* – phospholipid methyltransferase and *CKI1* – choline kinase) since these pathways are upregulated in *opi1Δ* cells under genotoxic stress (Figure 4D). Consequently, we deleted these genes in an *opi1Δ* background with the expectation that it would phenocopy the MMS resistance of an *opi1Δino2Δ* and *opi1Δino4Δ* strains. Surprisingly, all double mutant showed growth defects and increased MMS sensitivity (Supplementary Figure 7A and B). This negative genetic interaction between Opi1 and genes from PI and PC biosynthesis suggests that upregulation of one or more genes repressed by Opi1 in the absence of Ino1, Itr1, Opi3 and Cki1 negatively impacts the cell.

It is well established that the expression of *INO1* is positively regulated by Ino2-Ino4 and massively increases in the absence of Opi1 (Figure 3D and 4B) (Graves and Henry 2000; Santiago and Mamoun 2003; Jesch *et al*. 2005). To test if constitutive expression of *INO1* in cells lacking Opi1 is causing MMS sensitivity, we overexpressed *INO1* by cloning it under the control of the strong inducible *GAL1* promoter. Overexpression of *INO1* from the *GAL1* promoter rescues the inositol auxotrophy of an *ino1Δ* strain (Supplementary Figure 7C) but does not show sensitivity to MMS (Supplementary Figure 7D), which is an indication that increased levels of inositol do not affect the DDR.

### Inositol pyrophosphates counteract the effects of Opi1 on gene expression during genotoxic stress

In budding yeast, Kcs1 catalyzes the synthesis of inositol pyrophosphates. The inositol polyphosphates IP5 and IP6, which are generated by Ipk2 and Ipk1, respectively, serve as substrates for Kcs1, which catalyzes the pyrophosphorylation of the C5 α-phosphate to form 5-PP-IP4 and 5-PP-IP5, respectively (Saiardi *et al*. 1999). Miriam Greenberg’s group has proposed that inositol pyrophosphates produced in a Kcs1-dependent manner regulate the expression of *INO1* (Ye *et al*. 2013). The hypothesis suggests that the inositol pyrophosphate 5-PP-IP4 may play a role in recruiting transcriptional activators, such as Ino2-Ino4, to the promoter region of *INO1*. This suggests that deletion of Kcs1, involved in the synthesis of inositol pyrophosphates, could lead to a reduction in the expression of genes that are upregulated by Ino2-Ino4 in cells lacking Opi1. We consequently conducted a genetic rescue experiment by deleting Kcs1 in an *opi1Δ* strain to determine whether it would ameliorate the MMS sensitivity observed in the absence of Opi1. Remarkably, Kcs1 deletion significantly rescues the MMS sensitivity observed in the *opi1Δ* strain (Figure 5B). Additionally, Kcs1 deletion also rescue the G1/S arrest observed in *opi1Δ* cells during recover from MMS treatment (Figure 5C). These findings might indicate that in the presence of genotoxic stress induced by MMS, Opi1 functions in opposition to inositol pyrophosphates to facilitate the modulation of gene expression in an Ino2-Ino4 dependent manner. In agreement with that, cells lacking the Siw14 phosphatase that counteracts Kcs1-dependent PP-IPs synthesis have a mild sensitivity to MMS (Supplementary Figure 8A). It has been shown that *siw14Δ* mutants have increased levels of 5-PP-IP_5_ (Steidle *et al*. 2016).

To assess the impact of elevated inositol pyrophosphate levels on MMS sensitivity, we constructed a strain in which *KCS1* was placed under the control of the *GAL1* promoter. Our data demonstrate that induction with galactose results in elevated levels of Kcs1 mRNA and protein (see Supplementary Figure 8B and C). Remarkably, wild-type cells overexpressing Kcs1 phenocopy an *opi1Δ* strain with cell growth delay and sensitivity to MMS (Figure 5D). Taken together, our results suggest that increased PP– IPs levels due to the lack of the transcriptional repressor Opi1 leads to impaired capacity to cope with genotoxic stress induced by MMS.

Intriguingly, while the deletion of Ino2-Ino4 rescues the sensitivity to other genotoxic agents (Figure 5A), the deletion of Kcs1 in *opi1Δ* cells instead increase sensitivity to zeocin/bleomycin and 4NQO (Figure 5E). Furthermore, *opi1Δkcs1Δ* cells show sensitivity to HU and CPT (Supplementary Figure 8D). These findings suggest a complex interplay between Opi1 and inositol pyrophosphates in the cellular response to different sources of genotoxic stress. The specific mechanisms underlying the distinct responses to genotoxic agents warrant further investigation.

### Mitochondria function and genome stability

Remarkably, our RNA-seq data, combined with gene ontology analysis demonstrates a significant enrichment of genes with mitochondrial localization that show significant upregulation or downregulation (log_2_ fold change ≥ 0.8 or ≤ 0.8 respectively) in response to MMS treatment in *opi1Δ* cells (Figure 6A). Of note, it has been shown that the lack of Opi1 causes a decrease in mitochondrial membrane cardiolipin content leading to respiratory defects and a petite-negative phenotype (*pet^-^*) due to the inability to survive in the presence of damaged mitochondrial DNA (mtDNA) (Dunn *et al*. 2006; Luévano-Martínez *et al*. 2013). Interestingly, the expression of Opi1 is upregulated in cells lacking mitochondrial DNA (rho^0^), emphasizing the critical role of Opi1 in managing mitochondrial dysfunction (Singh *et al*. 2004). Our results confirm the sensitivity of Opi1-deficient cells to ethidium bromide, a known inducer of mtDNA damage, and we demonstrate that deletion of Ino2-Ino4 and Kcs1 rescues the *pet^-^* phenotype observed in *opi1Δ* cells (Figure 6B). Moreover, cells lacking Opi1 are sensitive to antimycin A, which can also be reverted by deletion of *INO2* and *INO4* (Supplementary Figure 9). Antimycin A, an inhibitor of complex III of the electron transport chain, can induce oxidative damage to mtDNA by increasing ROS leakage (Doudican *et al*. 2005).

Together, these data strongly suggest that constitutive activation of Ino2-Ino4 is also involved in the mitochondrial dysfunction probably due to mtDNA instability. Therefore, we hypothesize that the sensitivity of *opi1Δ* cells to MMS is more likely due to increased mtDNA instability rather than direct damage to nuclear DNA, as MMS is also known to induce damage to mtDNA (Kitanovic *et al*. 2009). Notably, the combined treatment of MMS and ethidium bromide did not result in an additive or synergistic increase in the sensitivity of *opi1Δ* cells, suggesting that the MMS sensitivity of these cells may be attributed to increased mtDNA damage rather than direct nuclear DNA damage (Figure 6C).

To elucidate if the rescue of MMS sensitivity is related to a recovery of mitochondrial function we sought to investigate if *INO*2 deletion would also rescue the respiratory defect and MMS sensitivity of *opi1Δ* cells in the presence of a respiratory carbon source. Utilizing the OROBOROS system, we measured the rate of oxygen consumption in respiratory medium (YPEG; 2% glycerol and 1% ethanol) and found that deletion of Opi1 led to a reduction in mitochondrial respiration, which is consistent with a defect in oxidative phosphorylation. Interestingly, the respiratory defect in Opi1-deficient cells was fully restored upon the deletion of *INO2* (Figure 6D). Surprisingly, the *opi1Δkcs1Δ* strain showed a significant rescue of mitochondrial respiration (Figure 6E). It is wort noting that cells lacking Kcs1 are typically unable to survive in a non-fermentable carbon source medium due to the critical role of inositol pyrophosphates in mediating the diauxic shift through the modulation of Gcr1, a key transcription factor involve in glycolysis regulation (Szijgyarto *et al*. 2011). Furthermore, we tested if the deletion of Ino2-Ino4 would rescue the MMS sensitivity of *opi1Δ* cells in the presence of a respiratory carbon source (Figure 6F). In the presence of galactose, which is known to induce simultaneous respiration and fermentation (Fendt and Sauer 2010), deletion of Ino2 and Ino4 completely restored the MMS sensitivity of *opi1Δ* cells. Importantly, in the presence of glycerol there was a significant rescue of the MMS sensitivity, which indicates that constitutive expression of genes regulated by Ino2-Ino4 increases mtDNA instability in the presence of MMS. Therefore, we conclude that deregulation of expression of genes regulated by Ino2-Ino4 affects mitochondria function potentially leading to mtDNA instability.

Overall, our findings indicate that under genotoxic stress, Opi1 is important to the fine-tuning of gene expression, to ensure appropriate cellular responses. Future investigations into these mechanisms will deepen our understanding of how cells maintain homeostasis and cope with genotoxic stress.

## Discussion

Through transcriptomic and functional analyses, our study provides evidence supporting the role of the transcriptional repressor Opi1 as a crucial regulator of gene expression during genotoxic stress. Our findings indicate that the constitutive activation of the transcriptional activator Ino2-Ino4 is associated with increased genotoxin sensitivity, cell cycle defects and mitochondrial disfunction. Importantly, reduction of inositol pyrophosphate synthesis resulting from the deletion of Kcs1 can ameliorate the MMS induced defects observed in *opi1Δ* cells. In contrast, the overexpression of Kcs1 in wild-type cells mimics the phenotype of cells lacking Opi1, suggesting that inositol pyrophosphates compete with Opi1 for the expression of genes in response to MMS treatment.

## Opi1 modulates different biological process during genotoxic stress

The regulation of multiple metabolic pathways by Opi1 poses a significant challenge in identifying the underlying cause of genotoxin sensitivity. In this study, we employed RNA-seq analyses to uncover Opi1’s modulation of diverse biological processes during genotoxic stress induced by MMS. Importantly, our findings underscore the pivotal role of Opi1 in the repression of phospholipid biosynthesis after genotoxic stress. It is worth noting that despite the involvement of Opi1 in regulating genes involved in phospholipid biosynthesis, the deletion of several genes related to this process did not rescue the MMS sensitivity of *opi1Δ* cells. Surprisingly, Opi1 exhibited negative genetic interactions with genes involved in phosphatidylinositol (Ino1 and Itr1) and phosphatidylcholine synthesis (Opi3 and Cki1), as revealed by our Supplementary Figure 6A and D. One explanation is that impairment of PC synthesis leading to an unbalance of phosphatidylethanolamine/phosphatidylcholine ratio (PE/PC) causes ER stress (Thibault *et al*. 2012). Furthermore, accumulation of the PC metabolic intermediate phosphatidylmonomethylethanolamine (PMME) has been shown to exacerbate ER stress in yeast cells (Ishiwata-Kimata *et al*. 2022). Given the increased expression of genes upstream of Opi3, such as *CDS1*, *CHO1*, *PSD1*, *CHO2* (Table S8), it is conceivable that PMME levels are even higher in *opi1Δ* cells. Alternatively, the decrease in PC levels can result in reduced choline and phosphatidic acid (PA) levels through the action of Phospholipase D (*SPO14*). This reduction in PA, along with the upregulation of genes involved in PI and PC synthesis, may direct CDP-DAG towards the production of these phospholipids, thereby decreasing the synthesis of phosphatidylglycerol (PG) and, consequently, cardiolipin. Further investigation could help to clarify the specific mechanisms by which Opi1 and proteins involved in phospholipid biosynthesis are involved in the response to genotoxic stress.

Furthermore, our study revealed that multiple proteins involved in the transduction of mating signaling with cellular conjugation and invasive filamentous growth, are significantly downregulated during genotoxic stress in an Opi1-dependent manner (Figure 4D). Filamentous growth in yeast is triggered by nutrient limitation, and involves various pathways including MAPK, Ras/PKA, Snf1, and TOR. A comprehensive review of filamentous growth response is available in (Cullen and Sprague 2012). Interestingly, it has been shown that genotoxic stress induces filamentous growth in budding yeast (Jiang and Kang 2003) and in *Candida* species (Shi *et al*. 2007; Bravo Ruiz *et al*. 2020). Noteworthy, it has been proposed that *Saccharomyces cerevisiae* Opi1 regulates filamentous growth through the upregulation of Flo11, a GPI-anchored cell surface glycoprotein that determines colony morphology in yeast. Budding yeast cells lacking Opi1 exhibit impaired biofilm formation and invasive growth on agar plates, two conditions that are associated with the transition from yeast to filamentous form (Reynolds 2006). While we did not observe significant changes in Flo11 expression in *opi1Δ* cells, our RNA-seq analysis revealed significant upregulation of several genes that inhibit filamentous growth, especially after MMS treatment, including the Fus3 MAP kinase, the Nrg2 transcriptional repressor, and Mho1, a protein of unknown function. Interestingly, while Fus3 inhibits filamentous growth by causing degradation of the transcription factor Tec1 via phosphorylation (Bao *et al*. 2004; Chou *et al*. 2004), it promotes mating signaling by phosphorylating the CDK inhibitor Far1, leading to its association with Cln-Cdc28 (Peter *et al*. 1993; Tyers and Futcher 1993; Peter and Herskowitz 1994). Consequently, Far1 impedes the function of the Cln-Cdc28 complex in G1 phase, thus preventing cells from advancing from G1 to START (reviewed in (Bardwell 2005; Sieber *et al*. 2023). Our findings are intriguing as they provide evidence for the involvement of Opi1 in promoting the suppression of mating signaling under genotoxic stress conditions. To begin with, *FAR1* and *FUS3* expression are downregulated in an Opi1-dependent manner (Figure 4E). Moreover, we observed an increase in the percentage of cells in G1 phase following a 2-hour MMS treatment (Figure 2D and E). Furthermore, upon release from MMS treatment, *opi1Δ* cells exhibited a delayed G1 to S-phase transition, indicating that mating signaling is hyperactivated (Figure 2E).

It is important to note that all of our results were obtained using BY4741, which is known to be a filamentation-deficient deficient yeast strain. Thus, any phenotypes related to filamentous growth, such as biofilm formation and invasive growth on agar, were not observable in our study.

### Inositol pyrophosphates generated by Kcs1 dampen Opi1-mediated gene expression during genotoxic stress

Our findings reveal that the MMS sensitivity and cell cycle defects observed in *opi1Δ* cells can be rescued by deletion of the inositol pyrophosphate kinase Kcs1 (Figure 5B and C), which provides compelling evidence that PP-IPs synthesis in the absence of Opi1 is detrimental for cells under genotoxic stress induced by MMS. It has been previously shown that *KCS1* possesses UAS_INO_ sequences in its promoter region and that its expression is upregulated when inositol concentration increases (Wimalarathna *et al*. 2011). Additionally, the inositol polyphosphate multikinase Ipk2 (*ARG82*) that catalyzes the synthesis of IP_4_ and IP_5_ also contains UAS_INO_ sequences in its promoter region, and its expression is dependent on Ino2-Ino4 (Wimalarathna *et al*. 2011). However, we did not observe changes in K*CS1* and *ARG82* expression levels in our dataset for *opi1Δ* cells (Table S4, S6 and S8), but it is plausible that PP-IPs synthesis is elevated in these cells, as previous studies have demonstrated an increase in Kcs1 protein levels in the absence of Opi1 (Ye *et al*. 2013). It is interesting to note that while *INO1* overexpression does not sensitize cells to MMS, that of *KCS1* does (Supplementary Figure 6C and Figure 5D respectively), suggesting that *INO1* overexpression does not lead to an increase in PP-IPs synthesis, which might indicate that there is a rate-limiting step reaction downstream of inositol-3-phophate synthesis restraining PP-IPs concentration in the cell.

How does inositol pyrophosphates affect the DDR leading to MMS sensitivity in the absence of Opi1? In budding yeast, PP-IPs are synthetized from the inositol polyphosphates IP_5_ and IP_6_ in a reaction catalyzed by the PP-IPs kinases Kcs1 and Vip1 (Saiardi *et al*. 1999; Mulugu *et al*. 2007; Lee *et al*. 2008). Kcs1 catalyzes the conversion of IP_5_ and IP_6_ to different isomers of PP-IPs such as 5-PP-IP_4_ and 5-PP-IP_5_ (Saiardi *et al*. 1999). It is established that PP-IPs modulate protein function either through non-enzymatic pyrophosphorylation of proteins, or allosteric regulation through its interaction with protein domains (Wilson *et al*. 2013). However, up until now, only a few targets have been characterized. Kcs1 and its mammalian orthologs IP6Ks have been implicated in several biological processes such as cell cycle progression, DNA repair, telomeric length, nutrient signaling, environmental stress response, polyphosphate metabolism among others (Saiardi *et al*. 2005; Auesukaree *et al*. 2005; Szijgyarto *et al*. 2011; Banfic *et al*. 2013, 2016; Worley *et al*. 2013; Wilson *et al*. 2013). In light of these considerations, our findings suggest a potential role for inositol pyrophosphates in modulating the function of Opi1-dependent genes during genotoxic stress induced by MMS. One plausible explanation is the previous proposal that PP-IPs generated by Kcs1 are important to induce *INO1* expression, indicating their potential involvement in recruiting Ino2-Ino4 to UAS_INO_ regions (Ye *et al*. 2013). However, this hypothesis requires further investigation, as there is currently no evidence suggesting that inositol pyrophosphates (PP-IPs) directly target Ino2-Ino4 for either allosteric regulation or pyrophosphorylation. It is also important to note that while Kcs1 deletion rescues MMS sensitivity of *opi1Δ* cells, it shows a synergistic increase in sensitivity to other genotoxins (Figure 5E). It has been shown that Kcs1 is required for DNA hyperrecombination phenotype in yeast cells with defects in protein kinase C1 (Pkc1) (Luo *et al*. 2002). Moreover, it was shown in mammalian cells that the orthologous of Kcs1 (IP6K1/2/3) are important for homologous recombination repair (Jadav *et al*. 2013). Hence, it is important to conduct future investigations to examine the potential cooperative role between Opi1 and Kcs1 in dealing with specific types of DNA lesions and their impact on DNA repair pathways. Furthermore, unraveling the underlying mechanisms behind the rescue of MMS sensitivity by Kcs1, as well as its promotion of sensitivity to other genotoxins, warrants thorough exploration. By looking into these aspects, we can gain a deeper understanding of the role of Opi1 and Kcs1 for cellular responses to genotoxic stress.

### Genotoxin sensitivity in *opi1Δ* is associated with mitochondrial DNA damage

Deletion of the transcriptional repressor Opi1 can induce pleiotropic effects in the cell. Therefore, we conducted a careful analysis to rule out the possibility that the genotoxin sensitivity of *opi1Δ* cells were due to a deregulation of other metabolic pathways not directly associated with the DDR. First, we ruled out the possibility that the heightened sensitivity of *opi1Δ* cells to MMS is linked to deregulation of the environmental stress response (ESR), as exposure to heat shock, osmotic stress, and oxidative stress did not result in increased sensitivity (Figure 1D). Besides that, tunicamycin treatment did not sensitize *opi1Δ* cells (Figure 1D), which is in line with previous evidence that the absence of Opi1 causes an expansion of the ER membrane, alleviating the ER stress induced by DTT or tunicamycin in an UPR independent manner (Schuck *et al*. 2009). Furthermore, it was previously shown that cells lacking Opi1 possess short telomeres (Askree *et al*. 2004). In contrast, cells lacking Kcs1 have longer telomeres (York *et al*. 2005). This finding could indicate that deletion of Kcs1 rescues MMS sensitivity due to restoration of telomere length. However, this is probably not the case as it was shown that telomere length does not affect cellular fitness nor yeast sensitivity to DNA damage (Harari *et al*. 2017).

MMS can also damage mitochondrial DNA leading to direct inhibition of the respiratory chain with increased ROS leakage, oxidative inactivation of glycolytic enzymes and cell cycle arrest (Kitanovic *et al*. 2009). Importantly, cells lacking Opi1 exhibit a decrease in mitochondrial cardiolipin content and are unable to survive in the absence of mtDNA (pet^-^ phenotype). The authors of the manuscript proposed that overproduction of inositol may impact cardiolipin content (Luévano-Martínez *et al*. 2013). Here, we have discovered that the sensitivity of cells lacking Opi1 to MMS may be linked to mitochondrial DNA instability. This conclusion is supported by two observations: a) Deletion of Ino2-Ino4 genes rescues the pet^-^ phenotype, antimycin A hypersensitivity, and respiratory capacity of cells lacking Opi1; b) The combined treatment of MMS and ethidium bromide does not increase the sensitivity of *opi1Δ* cells. It is worth noting that a previous study has demonstrated that cells lacking mitochondrial DNA (mtDNA) exhibit defects in the G1 to S-phase progression (Crider *et al*. 2012). Therefore, it would be important to investigate whether Opi1 is involved in a mitochondria-to-nucleus retrograde signaling pathway that promotes the transition from G1 to S-phase.

Furthermore, we observed that there is a significant enrichment of mitochondrial proteins among the genes that are differentially expressed in *opi1Δ* cells following MMS treatment (Figure 6A). For instance, we observed upregulation of Psd1 in *opi1Δ* cells upon genotoxic stress, as depicted in Figure 4D and Table S6-S7. Psd1 is an enzyme located in the mitochondrial inner membrane that converts phosphatidylserine to phosphatidylethanolamine and plays a crucial role in regulating mitochondrial fusion and morphology (Chan and McQuibban 2012). In contrast, Aim17, a protein with a mitochondrial localization and unknown function, was found to be downregulated in *opi1Δ* cells upon genotoxic stress as shown in Supplementary Figure 5B and Table S6-S7. Although limited information is available regarding the precise role of Aim17 in mitochondrial homeostasis, it seems to be important for mitochondrial DNA integrity (Hess *et al*. 2009). Considering that Psd1 and Aim17 are among the top hits of differentially expressed genes in our screening (Table S6), it would be valuable to explore in future investigations the specific contributions of these genes to the observed genotoxin sensitivity and mitochondrial dysfunction in *opi1Δ* cells.

### A working model for Opi1 function during genotoxic stress

We propose a model in which Opi1 functions as a crucial sensor of genotoxic stress in budding yeast (Figure 7). Upon genotoxic stress, Opi1 translocates to the nucleus and regulates the expression of multiple genes either directly or indirectly. Specifically, during MMS-induced genotoxic stress, Opi1 represses phospholipid biosynthesis and MAPK signaling, potentially suppressing cellular conjugation with mating while promoting invasive filamentous growth. Additionally, Opi1 fosters the expression of genes involved in amino acid metabolism and sulfate assimilation. In cells lacking Opi1, the constitutive activation of Ino2-Ino4 promotes gene transcription, which is further potentiated by inositol pyrophosphates produced by Kcs1. Consequently, the deletion of Opi1 results in pleiotropic defects affecting cell cycle progression and mitochondrial function, ultimately reducing cellular fitness. Understanding the interconnections between these processes and determining their relative contributions to MMS resistance will be crucial for future investigations.

**Figure 7.**
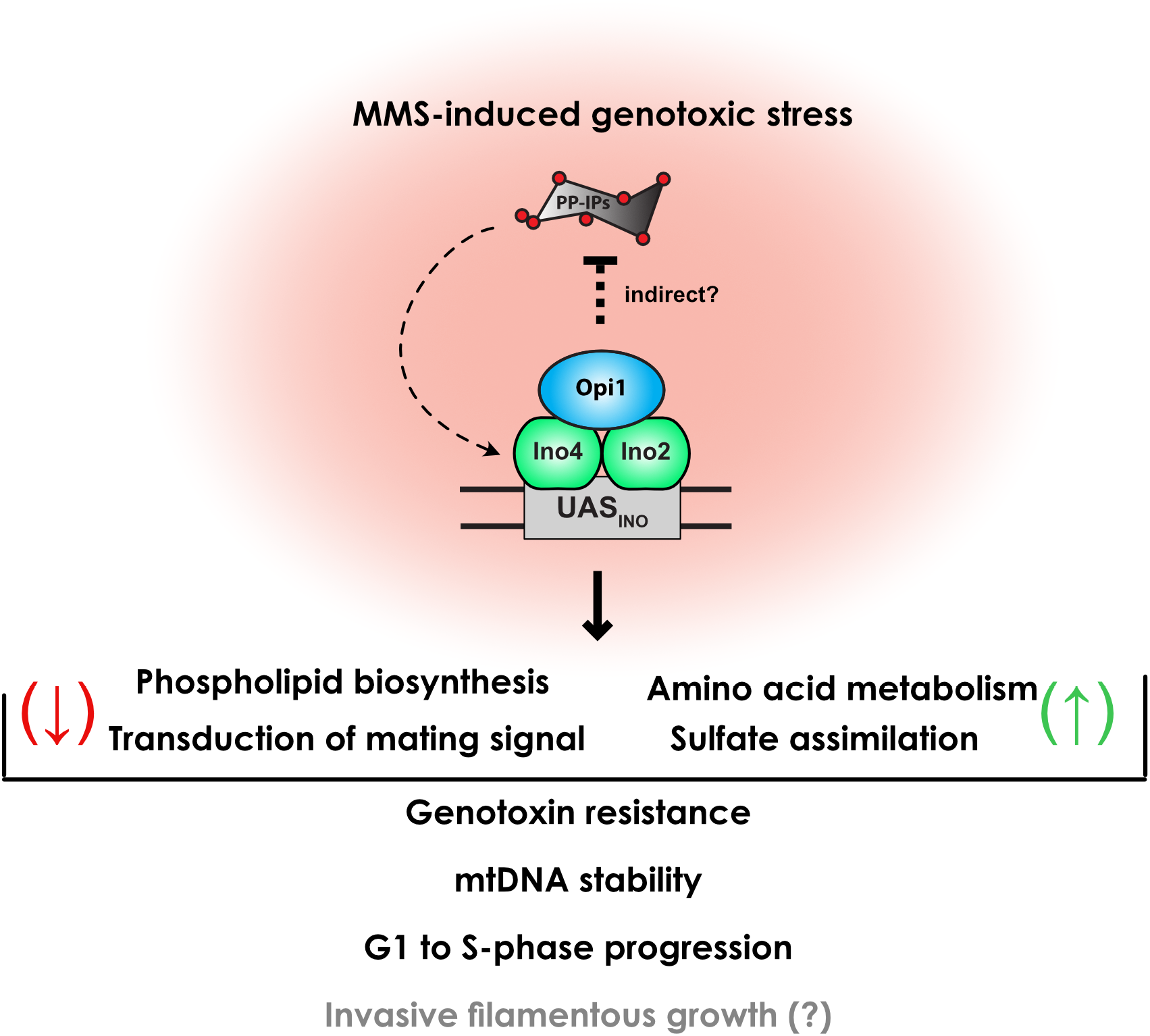
Model summarizing the Opi1 role as a key sensor of genotoxic stress in budding yeast. Briefly, upon genotoxic stress induced by MMS, Opi1 physically interacts with the Ino2-Ino4 transcriptional activator and counteracts inositol pyrophosphates, thereby modulating the expression of several genes. Opi1 plays an important role in repressing phospholipid biosynthesis and transduction of mating signaling, while promoting the expression of genes involved in amino acid metabolism and sulfate assimilation. The coordinated modulation of these genes contributes to genotoxin resistance by preventing mitochondrial DNA instability, delaying G1 to S-phase progression, and potentially promoting invasive filamentous growth. See discussion for more details.

In conclusion, this study provides insights into the intricate molecular mechanisms underlying Opi1’s role in cellular response to genotoxic stress. The significance of these findings is magnified by the presence of Opi1 in various pathogenic fungal species, which have become a global concern due to their widespread distribution and increasing antifungal resistance. Importantly, as Opi1 is absent in mammals, our results highlight its potential as a target for the development of more effective treatments against pathogenic fungal infections.

## Acknowledgements

We thank Jacilene Barbosa and Natalia Bromberg for technical support. Dr. Aparecida Sadae Tanaka and Dr. Maria Luiza Vilela Oliva for kindly allowing access to their laboratory and equipment. Dr. Francisco Bastos de Oliveira and Dr. Gustavo Monteiro Silva for comments and suggestions, Dr. Francisco Bastos de Oliveira, Dr Marcus Bustamante Smolka and Dr. Luis Eduardo Soares Netto for donating plasmid and yeast strains.

## Author contributions

J.R.C. conceived the project and wrote the paper. J.R.C., G.M.P., E.T-T., M.R.P., R.R.F., R.J., designed the experiments, analyzed the data, and helped revised the manuscript. F.M.C. designed and performed experiments for measuring oxygen consumption and helped revised the manuscript. N.C.S-P and O.B. provided reagents, infrastructure and helped revised the manuscript.

## References

Askree S. H., T. Yehuda, S. Smolikov, R. Gurevich, J. Hawk, et al., 2004 A genome-wide screen for *Saccharomyces cerevisiae* deletion mutants that affect telomere length. Proc Natl Acad Sci U S A 101: 8658–63. https://doi.org/10.1073/pnas.0401263101

Auesukaree C., H. Tochio, M. Shirakawa, Y. Kaneko, and S. Harashima, 2005 Plc1p, Arg82p, and Kcs1p, enzymes involved in inositol pyrophosphate synthesis, are essential for phosphate regulation and polyphosphate accumulation in Saccharomyces cerevisiae. J Biol Chem 280: 25127–25133. https://doi.org/10.1074/jbc.M414579200

Balint A., T. Kim, D. Gallo, J. R. Cussiol, F. M. Bastos De Oliveira, et al., 2015 Assembly of Slx4 signaling complexes behind DNA replication forks. EMBO Journal 34: 2182–2197. https://doi.org/10.15252/embj.201591190

Banfic H., A. Bedalov, J. York, and D. Visnjic, 2013 Inositol pyrophosphates modulate S phase progression after pheromone-induced arrest in *Saccharomyces cerevisiae*. J Biol Chem 288: 1717–1725. https://doi.org/10.1074/jbc.M112.412288

Banfic H., V. Crljen, L.-S. Vesna, V. Dembitz, H. Lalic, et al., 2016 Inositol pyrophosphates modulate cell cycle independently of alteration in telomere length. Adv Biological Regul 60: 22–28. https://doi.org/10.1016/j.jbior.2015.09.003

Bao M. Z., M. A. Schwartz, G. T. Cantin, J. R. Yates, and H. D. Madhani, 2004 Pheromone-dependent destruction of the Tec1 transcription factor is required for MAP kinase signaling specificity in yeast. Cell 119: 991–1000. https://doi.org/10.1016/j.cell.2004.11.052

Bardwell L., 2005 A walk-through of the yeast mating pheromone response pathway. Peptides (N.Y.) 26: 339–50. https://doi.org/10.1016/j.peptides.2004.10.002

Bastos de Oliveira F. M., D. Kim, J. R. Cussiol, J. Das, M. Jeong, et al., 2015 Phosphoproteomics reveals distinct modes of Mec1/ATR signaling during DNA replication. Mol Cell 57. https://doi.org/10.1016/j.molcel.2015.01.043

Beranek D. T., 1990 Distribution of methyl and ethyl adducts following alkylation with monofunctional alkylating agents. Mutat Res 231: 11–30. https://doi.org/10.1016/0027-5107(90)90173-2

Branzei D., and M. Foiani, 2006 The Rad53 signal transduction pathway: Replication fork stabilization, DNA repair, and adaptation. Exp. Cell Res. 312: 2654–2659. https://doi.org/10.1016/j.yexcr.2006.06.012

Branzei D., and M. Foiani, 2010 Maintaining genome stability at the replication fork. Nat. Rev. Mol. Cell Biol. 11: 208–219. https://doi.org/10.1038/nrm2852

Bravo Ruiz G., Z. K. Ross, N. A. R. Gow, and A. Lorenz, 2020 Pseudohyphal growth of the emerging pathogen *Candida auris* is triggered by genotoxic stress through the S phase checkpoint. mSphere 5. https://doi.org/10.1128/mSphere.00151-20

Brickner J. H., and P. Walter, 2004 Gene recruitment of the activated *INO1* locus to the nuclear membrane. PLoS Biol 2: e342. https://doi.org/10.1371/journal.pbio.0020342

Chan E. Y. L., and G. A. McQuibban, 2012 Phosphatidylserine decarboxylase 1 (Psd1) promotes mitochondrial fusion by regulating the biophysical properties of the mitochondrial membrane and alternative topogenesis of mitochondrial genome maintenance protein 1 (Mgm1). J Biol Chem 287: 40131–9. https://doi.org/10.1074/jbc.M112.399428

Chou S., L. Huang, and H. Liu, 2004 Fus3-regulated Tec1 degradation through SCFCdc4 determines MAPK signaling specificity during mating in yeast. Cell 119: 981–90. https://doi.org/10.1016/j.cell.2004.11.053

Chumnanpuen P., I. Nookaew, and J. Nielsen, 2013 Integrated analysis, transcriptome-lipidome, reveals the effects of INO-level (INO2 and INO4) on lipid metabolism in yeast. BMC Syst Biol 7: S7. https://doi.org/10.1186/1752-0509-7-S3-S7

Craven R. J., and T. D. Petes, 2001 The *Saccharomyces cerevisiae* suppressor of choline sensitivity (*SCS2*) gene is a multicopy Suppressor of mec1 telomeric silencing defects. Genetics 158: 145–54. https://doi.org/10.1093/genetics/158.1.145

Crider D. G. G.-R. L. J, P. Srivastava, P.-R. Leonardo, K. Upadhyaya, et al., 2012 Rad53 is essential for a mitochondrial {DNA} inheritance checkpoint regulating G1 to S progression. J. Cell Biol. 198: 793–798. https://doi.org/10.1083/jcb.201205193

Cullen P. J., and G. F. Sprague, 2012 The regulation of filamentous growth in yeast. Genetics 190: 23–49. https://doi.org/10.1534/genetics.111.127456

Cussiol J. R. R., B. L. Soares, and F. M. B. de Oliveira, 2020 From yeast to humans: Understanding the biology of DNA damage response (DDR) kinases. Genet Mol Biol 43. https://doi.org/10.1590/1678-4685-gmb-2019-0071

Dobin A., C. A. Davis, F. Schlesinger, J. Drenkow, C. Zaleski, et al., 2013 STAR: ultrafast universal RNA-seq aligner. Bioinformatics 29: 15–21. https://doi.org/10.1093/bioinformatics/bts635

Doudican N. A., B. Song, G. S. Shadel, and P. W. Doetsch, 2005 Oxidative DNA damage causes mitochondrial genomic instability in *Saccharomyces cerevisiae*. Mol Cell Biol 25: 5196–204. https://doi.org/10.1128/MCB.25.12.5196-5204.2005

Downs J. A., N. F. Lowndes, and S. P. Jackson, 2000 A role for *Saccharomyces cerevisiae* histone H2A in DNA repair. Nature 408: 1001–4. https://doi.org/10.1038/35050000

Dunn C. D., M. S. Lee, F. A. Spencer, and R. E. Jensen, 2006 A genomewide screen for petite-negative yeast strains yields a new subunit of the i-AAA protease complex. Mol Biol Cell 17: 213–26. https://doi.org/10.1091/mbc.e05-06-0585

Fendt S.-M., and U. Sauer, 2010 Transcriptional regulation of respiration in yeast metabolizing differently repressive carbon substrates. BMC Syst Biol 4: 12. https://doi.org/10.1186/1752-0509-4-12

Fernandez-Capetillo O., A. Lee, M. Nussenzweig, and A. Nussenzweig, 2004 H2AX: the histone guardian of the genome. DNA Repair (Amst) 3: 959–967. https://doi.org/10.1016/j.dnarep.2004.03.024

Ferrari E., C. Bruhn, M. Peretti, C. Cassani, W. V. Carotenuto, et al., 2017 PP2A controls genome integrity by integrating nutrient-sensing and metabolic pathways with the DNA Damage Response. Mol Cell 67: 266–281.e4. https://doi.org/10.1016/j.molcel.2017.05.027

Gasch A. P., M. Huang, S. Metzner, D. Botstein, S. J. Elledge, et al., 2001 Genomic expression responses to DNA-damaging agents and the regulatory role of the yeast ATR homolog Mec1p. Mol Biol Cell 12: 2987–3003. https://doi.org/10.1091/mbc.12.10.2987

Gaspar M. L., Y.-F. Chang, S. A. Jesch, M. Aregullin, and S. A. Henry, 2017 Interaction between repressor Opi1p and ER membrane protein Scs2p facilitates transit of phosphatidic acid from the ER to mitochondria and is essential for *INO1* gene expression in the presence of choline. J Biol Chem 292: 18713–18728. https://doi.org/10.1074/jbc.M117.809970

Gietz D., S. A. Jean, R. A. Woods, and R. H. Schiestl, 1992 Improved method for high efficiency transformation of intact yeast cells. Nucleic Acids Res 20: 1425.

Gietz R. D., and R. A. Woods, 2006 Yeast transformation by the LiAc/SS Carrier DNA/PEG method. Methods Mol Biol 313: 107–120. https://doi.org/10.1385/1-59259-958-3:107

Graves J. A., and S. A. Henry, 2000 Regulation of the yeast INO1 gene: The products of the INO2, INO4 and OPI1 regulatory genes are not required for repression in response to inositol. Genetics 154: 1485–1495.

Haber J. E., and W. Y. Leung, 1996 Lack of chromosome territoriality in yeast: promiscuous rejoining of broken chromosome ends. Proc Natl Acad Sci U S A 93: 13949–54. https://doi.org/10.1073/pnas.93.24.13949

Hanawalt P. C., 2015 Historical perspective on the DNA damage response. DNA Repair (Amst) 36: 2–7. https://doi.org/10.1016/j.dnarep.2015.10.001

Harari Y., S. Zadok-Laviel, and M. Kupiec, 2017 Long telomeres do not affect cellular fitness in yeast. mBio 8: 1–9. https://doi.org/10.1128/mBio.01314-17

Hess D. C., C. L. Myers, C. Huttenhower, M. A. Hibbs, A. P. Hayes, et al., 2009 Computationally driven, quantitative experiments discover genes required for mitochondrial biogenesis. PLoS Genet 5: e1000407. https://doi.org/10.1371/journal.pgen.1000407

Heyken W.-T. T., A. Repenning, J. Kumme, and H.-J. J. Schüller, 2005 Constitutive expression of yeast phospholipid biosynthetic genes by variants of Ino2 activator defective for interaction with Opi1 repressor. Mol. Microbiol. 56: 696–707. https://doi.org/10.1111/j.1365-2958.2004.04499.x

Hirsch J. P., and S. A. Henry, 1986 Expression of the *Saccharomyces cerevisiae* inositol-1-phosphate synthase (INO1) gene is regulated by factors that affect phospholipid synthesis. Mol Cell Biol 6: 3320–8. https://doi.org/10.1128/mcb.6.10.3320-3328.1986

Hofbauer H. F., M. Gecht, S. C. Fischer, A. Seybert, A. S. Frangakis, et al., 2018 The molecular recognition of phosphatidic acid by an amphipathic helix in Opi1. J Cell Biol 217: 3109– 3126. https://doi.org/10.1083/jcb.201802027

Hoppen J., A. Repenning, A. Albrecht, S. Geburtig, and H.-J. Schüller, 2005 Comparative analysis of promoter regions containing binding sites of the heterodimeric transcription factor Ino2/Ino4 involved in yeast phospholipid biosynthesis. Yeast 22: 601–613. https://doi.org/10.1002/yea.1209

Ishiwata-Kimata Y., Q. G. Le, and Y. Kimata, 2022 Induction and aggravation of the endoplasmic-reticulum stress by membrane-lipid metabolic intermediate Phosphatidyl-N-Monomethylethanolamine. Front Cell Dev Biol 9. https://doi.org/10.3389/fcell.2021.743018

Jablonowski C. M., J. R. Cussiol, S. Oberly, A. Yimit, A. Balint, et al., 2015 Termination of replication stress signaling via concerted action of the Slx4 scaffold and the PP4 phosphatase. Genetics 201. https://doi.org/10.1534/genetics.115.181479

Jackson S. P., and J. Bartek, 2009 The DNA-damage response in human biology and disease. Nature 461: 1071–1078.

Jadav R. S., M. V. L. Chanduri, S. Sengupta, and R. Bhandari, 2013 Inositol pyrophosphate synthesis by inositol hexakisphosphate kinase 1 is required for homologous recombination repair. J Biol Chem 288: 3312–21. https://doi.org/10.1074/jbc.M112.396556

Jäschke Y., J. Schwarz, D. Clausnitzer, C. Müller, and H.-J. Schüller, 2011 Pleiotropic corepressors Sin3 and Ssn6 interact with repressor Opi1 and negatively regulate transcription of genes required for phospholipid biosynthesis in the yeast *Saccharomyces cerevisiae*. Mol Genet Genomics 285: 91–100. https://doi.org/10.1007/s00438-010-0589-5

Jelinsky S. A., and L. D. Samson, 1999 Global response of *Saccharomyces cerevisiae* to an alkylating agent. Proc Natl Acad Sci U S A 96: 1486–91. https://doi.org/10.1073/pnas.96.4.1486

Jesch S. A., X. Zhao, M. T. Wells, and S. A. Henry, 2005 Genome-wide analysis reveals inositol, not choline, as the major effector of Ino2p-Ino4p and unfolded protein response target gene expression in yeast. J Biol Chem 280: 9106–18. https://doi.org/10.1074/jbc.M411770200

Jiang Y. W., and C. M. Kang, 2003 Induction of S. cerevisiae Filamentous Differentiation by Slowed DNA Synthesis Involves Mec1, Rad53 and Swe1 Checkpoint Proteins. Mol Biol Cell 14: 5116–5124. https://doi.org/10.1091/mbc.e03-06-0375

Kitanovic A., and S. Wölfl, 2006 Fructose-1,6-bisphosphatase mediates cellular responses to DNA damage and aging in *Saccharomyces cerevisiae*. Mutat Res 594: 135–47. https://doi.org/10.1016/j.mrfmmm.2005.08.005

Kitanovic A., T. Walther, M. O. Loret, J. Holzwarth, I. Kitanovic, et al., 2009 Metabolic response to MMS-mediated DNA damage in *Saccharomyces cerevisiae* is dependent on the glucose concentration in the medium. FEMS Yeast Res 9: 535–51. https://doi.org/10.1111/j.1567-1364.2009.00505.x

Kliewe F., M. Engelhardt, R. Aref, and H.-J. Schüller, 2017 Promoter recruitment of corepressors Sin3 and Cyc8 by activator proteins of the yeast Saccharomyces cerevisiae. Curr Genet 63: 739–750. https://doi.org/10.1007/s00294-017-0677-8

Lanz M. C., D. Dibitetto, and M. B. Smolka, 2019 DNA damage kinase signaling: checkpoint and repair at 30 years. EMBO J 38: e101801. https://doi.org/10.15252/embj.2019101801

Lanz M. C., K. Yugandhar, S. Gupta, E. J. Sanford, V. M. Faça, et al., 2021 In-depth and 3-dimensional exploration of the budding yeast phosphoproteome. EMBO Rep 22. https://doi.org/10.15252/embr.202051121

Lee Y. S., K. Huang, F. A. Quiocho, and E. K. O’Shea, 2008 Molecular basis of cyclin-CDK-CKI regulation by reversible binding of an inositol pyrophosphate. Nat Chem Biol 4: 25–32. https://doi.org/10.1038/nchembio.2007.52

Longtine M. S., A. McKenzie, D. J. Demarini, N. G. Shah, A. Wach, et al., 1998 Additional modules for versatile and economical PCR-based gene deletion and modification in Saccharomyces cerevisiae. Yeast 14: 953–961. https://doi.org/10.1002/(SICI)1097-0061(199807)14:10<953::AID-YEA293>3.0.CO;2-U

Love M. I., W. Huber, and S. Anders, 2014 Moderated estimation of fold change and dispersion for RNA-seq data with DESeq2. Genome Biol 15: 550. https://doi.org/10.1186/s13059-014-0550-8

Luévano-Martínez L. A., P. Appolinario, S. Miyamoto, S. Uribe-Carvajal, and A. J. Kowaltowski, 2013 Deletion of the transcriptional regulator opi1p decreases cardiolipin content and disrupts mitochondrial metabolism in *Saccharomyces cerevisiae*. Fungal Genet Biol 60: 150–8. https://doi.org/10.1016/j.fgb.2013.03.005

Luo H. R., A. Saiardi, H. Yu, E. Nagata, K. Ye, et al., 2002 Inositol pyrophosphates are required for DNA hyperrecombination in protein kinase C1 mutant yeast. Biochemistry 41: 2509– 2515. https://doi.org/10.1021/bi0118153

Mulugu S., W. Bai, P. C. Fridy, R. J. Bastidas, J. C. Otto, et al., 2007 A conserved family of enzymes that phosphorylate inositol hexakisphosphate. Science (1979) 316: 106–109. https://doi.org/10.1126/science.1139099

Pellicioli A., and M. Foiani, 2005 Signal transduction: how Rad53 kinase is activated. Curr. Biol. 15: R769–71. https://doi.org/10.1016/j.cub.2005.08.057

Peter M., A. Gartner, J. Horecka, G. Ammerer, and I. Herskowitz, 1993 FAR1 links the signal transduction pathway to the cell cycle machinery in yeast. Cell 73: 747–60. https://doi.org/10.1016/0092-8674(93)90254-n

Peter M., and I. Herskowitz, 1994 Direct inhibition of the yeast cyclin-dependent kinase Cdc28-Cln by Far1. Science 265: 1228–31. https://doi.org/10.1126/science.8066461

Reynolds T. B., 2006 The Opi1p Transcription Factor Affects Expression of *FLO11*, Mat Formation, and Invasive Growth in *Saccharomyces cerevisiae*. Eukaryot Cell 5: 1266–1275. https://doi.org/10.1128/EC.00022-06

Rogakou E. P., D. R. Pilch, A. H. Orr, V. S. Ivanova, and W. M. Bonner, 1998 DNA double-stranded breaks induce histone H2AX phosphorylation on serine 139. J Biol Chem 273: 5858–68. https://doi.org/10.1074/jbc.273.10.5858

Rothstein R. J., 1983 One-step gene disruption in yeast, pp. 202–211 in Methods Enzymol,.

Rothstein R., 1991 Targeting, disruption, replacement, and allele rescue: Integrative DNA transformation in yeast, pp. 281–301 in Methods Enzymol,.

Saiardi A., H. Erdjument-Bromage, A. M. Snowman, P. Tempst, and S. H. Snyder, 1999 Synthesis of diphosphoinositol pentakisphosphate by a newly identified family of higher inositol polyphosphate kinases. Curr Biol 9: 1323–6. https://doi.org/10.1016/s0960-9822(00)80055-x

Saiardi A., A. C. Resnick, A. M. Snowman, B. Wendland, and S. H. Snyder, 2005 Inositol pyrophosphates regulate cell death and telomere length through phosphoinositide 3-kinase-related protein kinases. Proc Natl Acad Sci U S A. https://doi.org/10.1073/pnas.0409322102

Salmon T. B., B. A. Evert, B. Song, and P. W. Doetsch, 2004 Biological consequences of oxidative stress-induced DNA damage in Saccharomyces cerevisiae. Nucleic Acids Res 32: 3712–23. https://doi.org/10.1093/nar/gkh696

Santiago T. C., and C. Ben Mamoun, 2003 Genome expression analysis in yeast reveals novel transcriptional regulation by inositol and choline and new regulatory functions for Opi1p, Ino2p, and Ino4p. J Biol Chem 278: 38723–30. https://doi.org/10.1074/jbc.M303008200

Schuck S., W. A. Prinz, K. S. Thorn, C. Voss, and P. Walter, 2009 Membrane expansion alleviates endoplasmic reticulum stress independently of the unfolded protein response. J Cell Biol 187: 525–36. https://doi.org/10.1083/jcb.200907074

Schüller H. J., A. Hahn, F. Tröster, A. Schütz, and E. Schweizer, 1992 Coordinate genetic control of yeast fatty acid synthase genes FAS1 and FAS2 by an upstream activation site common to genes involved in membrane lipid biosynthesis. EMBO J 11: 107–114. https://doi.org/10.1002/j.1460-2075.1992.tb05033.x

Shi Q.-M., Y.-M. Wang, X.-D. Zheng, R. Teck Ho Lee, and Y. Wang, 2007 Critical Role of DNA Checkpoints in Mediating Genotoxic-Stress–induced Filamentous Growth in *Candida albicans*. Mol Biol Cell 18: 815–826. https://doi.org/10.1091/mbc.e06-05-0442

Shirahige K., Y. Hori, K. Shiraishi, M. Yamashita, K. Takahashi, et al., 1998 Regulation of DNA-replication origins during cell-cycle progression. Nature 395: 618–621. https://doi.org/10.1038/27007

Sieber B., J. M. Coronas-Serna, and S. G. Martin, 2023 A focus on yeast mating: From pheromone signaling to cell-cell fusion. Semin Cell Dev Biol 133: 83–95. https://doi.org/10.1016/j.semcdb.2022.02.003

Simpson-Lavy K. J., A. Bronstein, M. Kupiec, and M. Johnston, 2015 Cross-Talk between Carbon Metabolism and the DNA Damage Response in *S. cerevisiae*. Cell Rep 12: 1865–75. https://doi.org/10.1016/j.celrep.2015.08.025

Singh K. K., A. K. Rasmussen, and L. J. Rasmussen, 2004 Genome-wide analysis of signal transducers and regulators of mitochondrial dysfunction in *Saccharomyces cerevisiae*. Ann N Y Acad Sci 1011: 284–98. https://doi.org/10.1007/978-3-662-41088-2_27

Steidle E. A., L. S. Chong, M. Wu, E. Crooke, D. Fiedler, et al., 2016 A Novel Inositol Pyrophosphate Phosphatase in *Saccharomyces cerevisiae*: Siw14 protein selectively cleaves the β-phosphate from 5-diphosphoinositol pentakisphosphate (5PP-IP5). J Biol Chem 291: 6772–83. https://doi.org/10.1074/jbc.M116.714907

Szijgyarto Z., A. Garedew, C. Azevedo, and A. Saiardi, 2011 Influence of inositol pyrophosphates on cellular energy dynamics. Science (1979) 334: 802–805. https://doi.org/10.1126/science.1211908

Szklarczyk D., A. L. Gable, D. Lyon, A. Junge, S. Wyder, et al., 2019 STRING v11: protein– protein association networks with increased coverage, supporting functional discovery in genome-wide experimental datasets. Nucleic Acids Res 47: D607–D613. https://doi.org/10.1093/nar/gky1131

Tercero J. A., and J. F. X. Diffley, 2001 Regulation of DNA replication fork progression through damaged DNA by the Mec1/Rad53 checkpoint. Nature 412: 553–557. https://doi.org/10.1038/35087607

Thibault G., G. Shui, W. Kim, G. C. McAlister, N. Ismail, et al., 2012 The membrane stress response buffers lethal effects of lipid disequilibrium by reprogramming the protein homeostasis network. Mol Cell 48: 16–27. https://doi.org/10.1016/j.molcel.2012.08.016

Thomas P. D., D. Ebert, A. Muruganujan, T. Mushayahama, L. Albou, et al., 2022 PANTHER: Making genome-scale phylogenetics accessible to all. Protein Science 31: 8–22. https://doi.org/10.1002/pro.4218

Tyers M., and B. Futcher, 1993 Far1 and Fus3 link the mating pheromone signal transduction pathway to three G1-phase Cdc28 kinase complexes. Mol Cell Biol 13: 5659–69. https://doi.org/10.1128/mcb.13.9.5659-5669.1993

Wagner C., M. Dietz, J. Wittmann, A. Albrecht, and H. J. Schüller, 2001 The negative regulator Opi1 of phospholipid biosynthesis in yeast contacts the pleiotropic repressor Sin3 and the transcriptional activator Ino2. Mol Microbiol 41: 155–66. https://doi.org/10.1046/j.1365-2958.2001.02495.x

Wang Y.-H., A. Hariharan, G. Bastianello, Y. Toyama, G. v. Shivashankar, et al., 2017 DNA damage causes rapid accumulation of phosphoinositides for ATR signaling. Nat Commun 8: 2118. https://doi.org/10.1038/s41467-017-01805-9

Wilson M. S. C., T. M. Livermore, and A. Saiardi, 2013 Inositol pyrophosphates: Between signalling and metabolism. Biochem J 452: 369–79. https://doi.org/10.1042/BJ20130118

Wimalarathna R., C.-H. Tsai, and C.-H. Shen, 2011 Transcriptional control of genes involved in yeast phospholipid biosynthesis. J Microbiol 49: 265–73. https://doi.org/10.1007/s12275-011-1130-1

Worley J., X. Luo, and A. P. Capaldi, 2013 Inositol pyrophosphates regulate cell growth and the environmental stress response by activating the HDAC Rpd3L. Cell Rep 3: 1476–82. https://doi.org/10.1016/j.celrep.2013.03.043

Wyatt M. D., and D. L. Pittman, 2006 Methylating agents and DNA repair responses: Methylated bases and sources of strand breaks. Chem Res Toxicol 19: 1580–94. https://doi.org/10.1021/tx060164e

Ye C., W. M. M. S. Bandara, and M. L. Greenberg, 2013 Regulation of inositol metabolism is fine-tuned by inositol pyrophosphates in *Saccharomyces cerevisiae*. J Biol Chem 288: 24898–908. https://doi.org/10.1074/jbc.M113.493353

Yi C., J. Tong, P. Lu, Y. Wang, J. Zhang, et al., 2017 Formation of a Snf1-Mec1-Atg1 module on mitochondria governs energy deprivation-induced autophagy by regulating mitochondrial respiration. Dev Cell 41: 59–71.e4. https://doi.org/10.1016/j.devcel.2017.03.007

York S. J., B. N. Armbruster, P. Greenwell, T. D. Petes, and J. D. York, 2005 Inositol diphosphate signaling regulates telomere length. J Biol Chem 280: 4264–9. https://doi.org/10.1074/jbc.M412070200

Zewail A., M. W. Xie, Y. Xing, L. Lin, P. F. Zhang, et al., 2003 Novel functions of the phosphatidylinositol metabolic pathway discovered by a chemical genomics screen with wortmannin. Proc Natl Acad Sci U S A 100: 3345–50. https://doi.org/10.1073/pnas.0530118100

Zhou C., A. E. H. Elia, M. L. Naylor, N. Dephoure, B. A. Ballif, et al., 2016 Profiling DNA damage-induced phosphorylation in budding yeast reveals diverse signaling networks. Proc Natl Acad Sci U S A 113: E3667–E3675. https://doi.org/10.1073/pnas.1602827113

Zulkifli M., J. K. Neff, S. A. Timbalia, N. M. Garza, Y. Chen, et al., 2020 Yeast homologs of human MCUR1 regulate mitochondrial proline metabolism. Nat Commun 11: 4866. https://doi.org/10.1038/s41467-020-18704-1

