## Supplementary Figures for "Opi1-mediated transcriptional modulation orchestrates genotoxic stress response in budding yeast"

#### Supplementary Figure 1

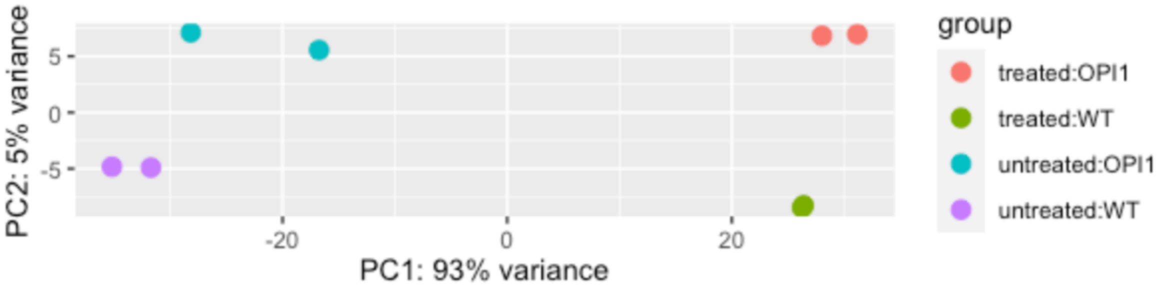

### Supplementary Figure 2

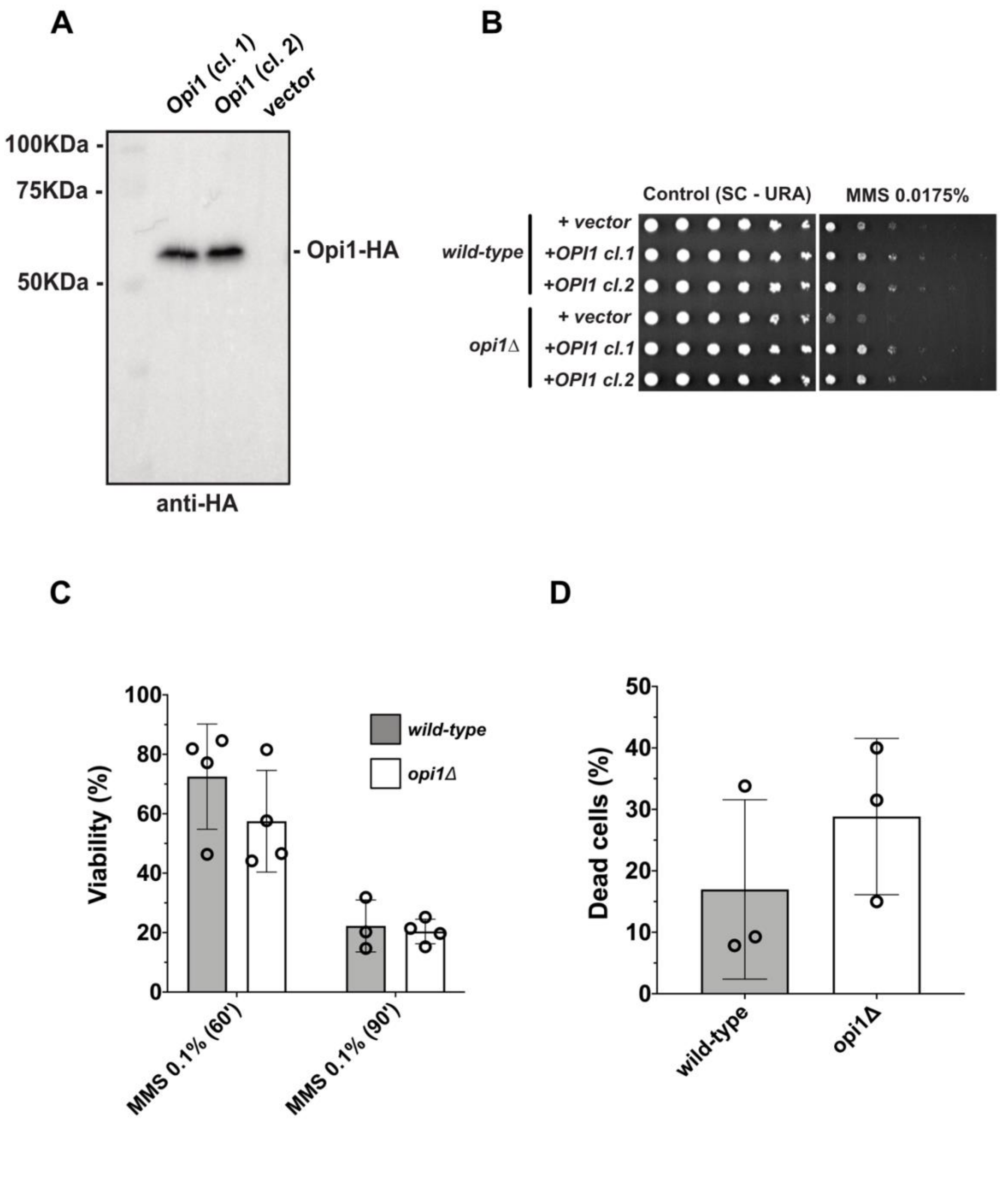

#### Supplementary Figure 3

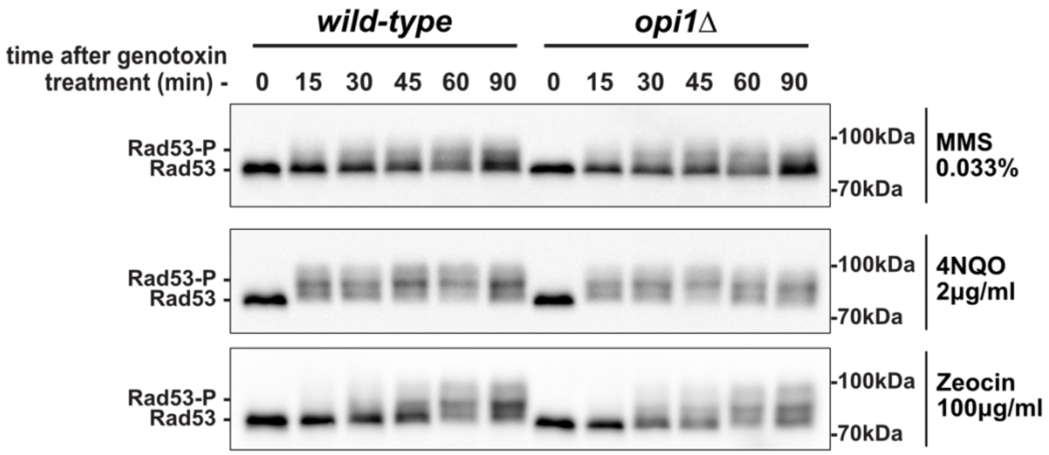

### Supplementary Figure 4

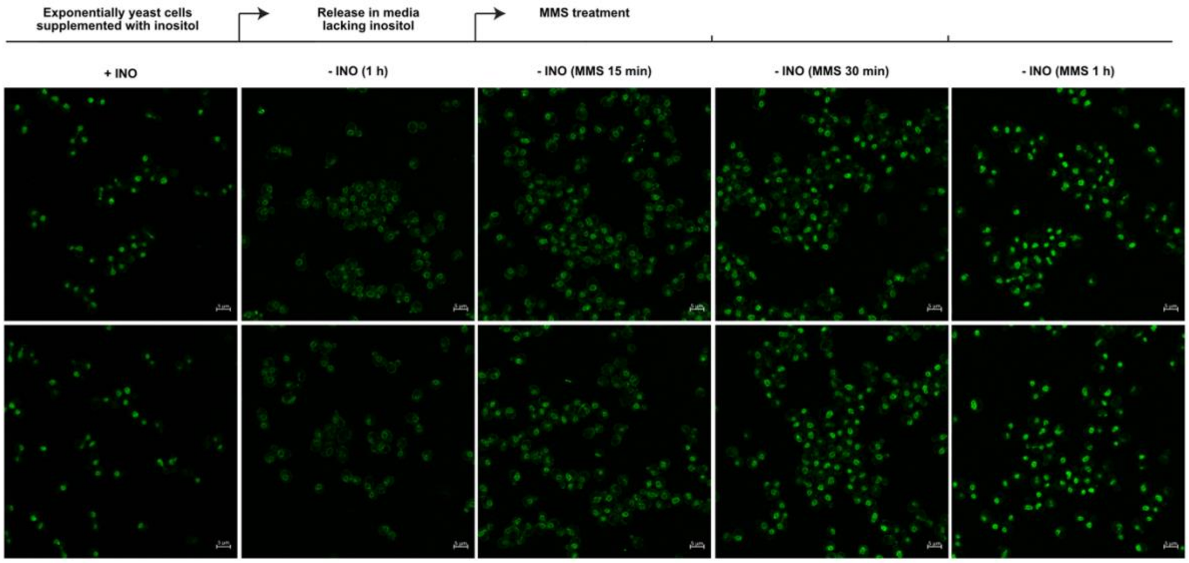

#### Supplementary Figure 5

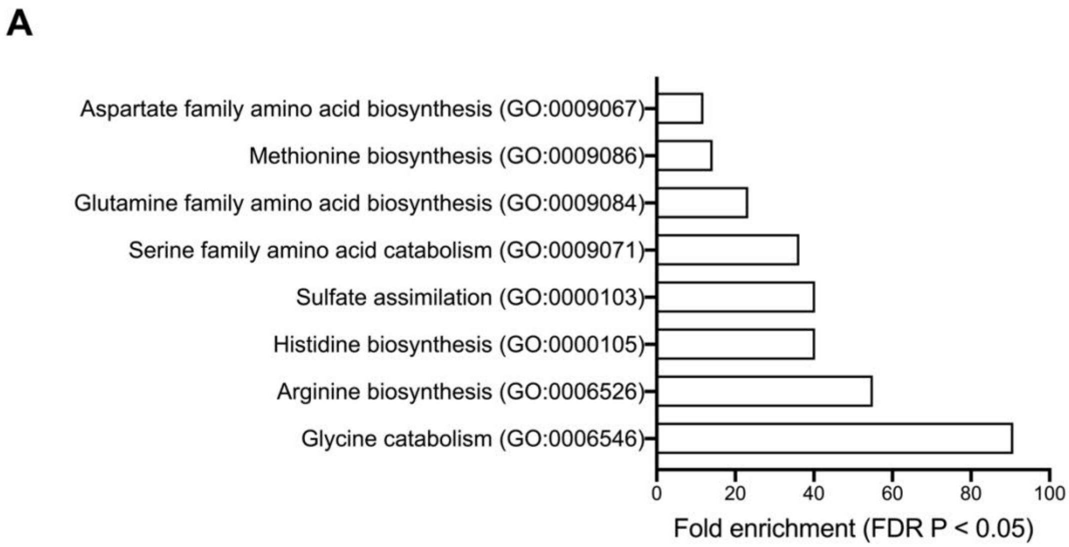


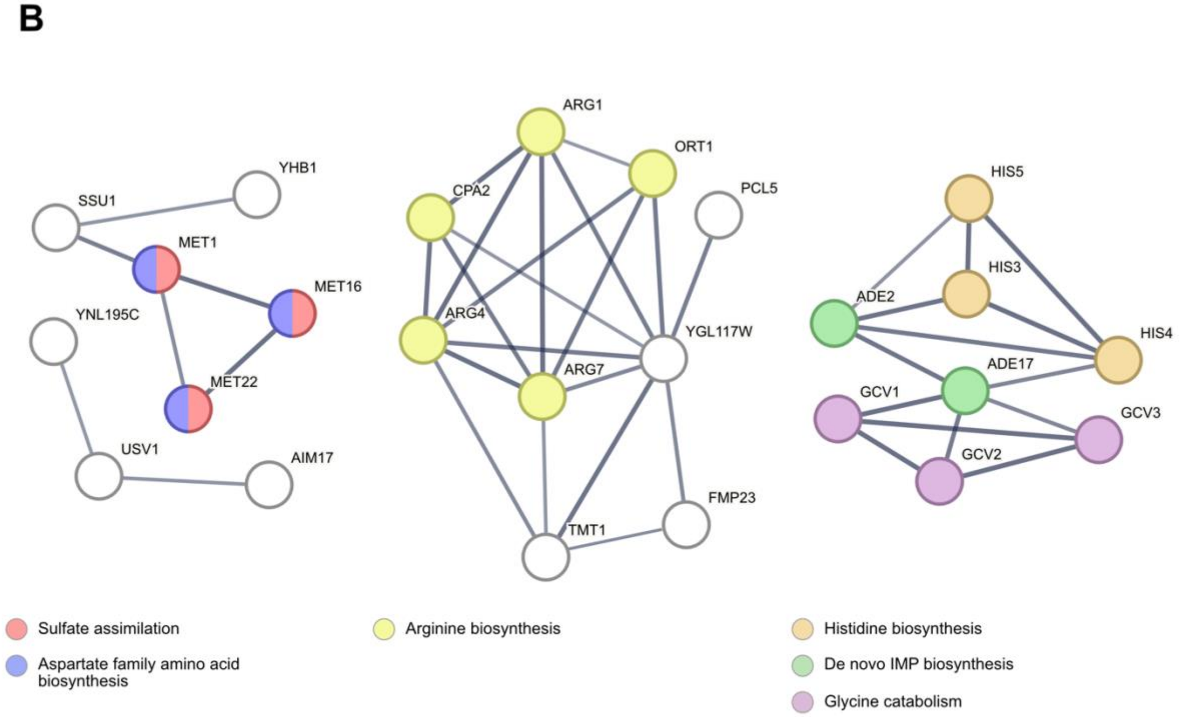

#### Supplementary Figure 6

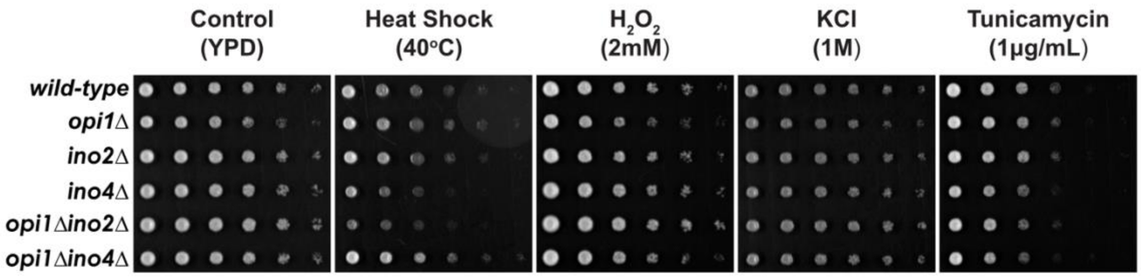

### Supplementary Figure 7

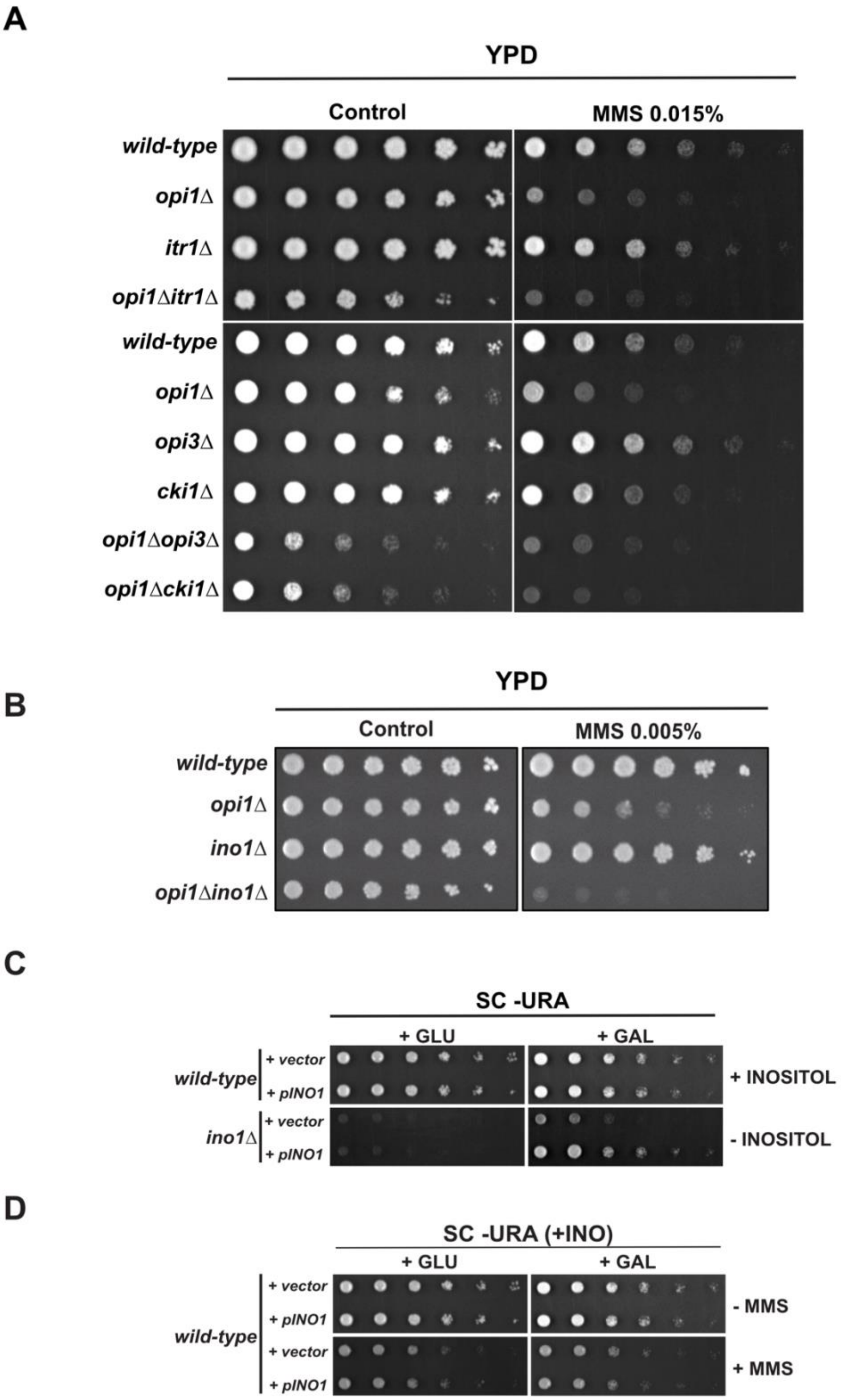

### Supplementary Figure 8

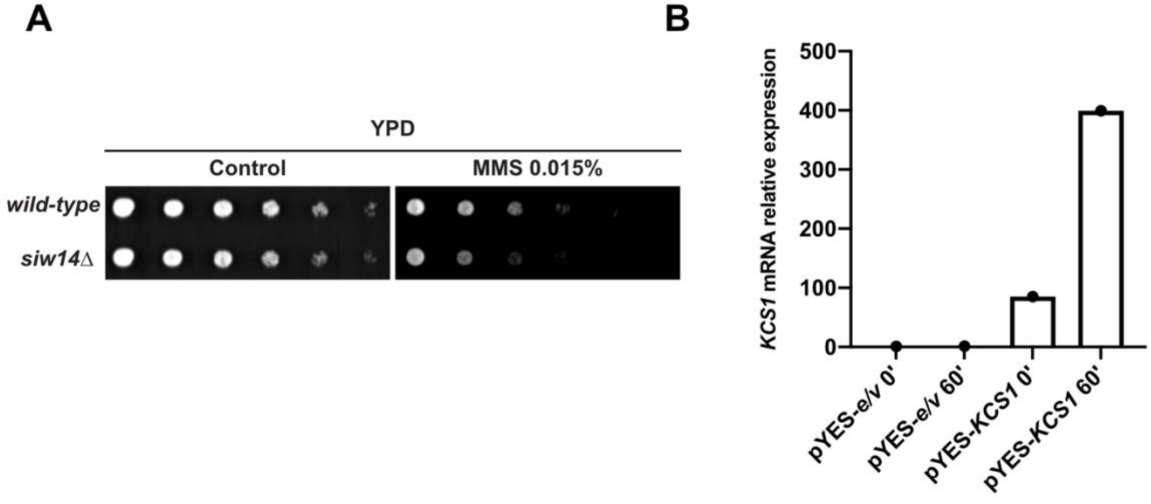


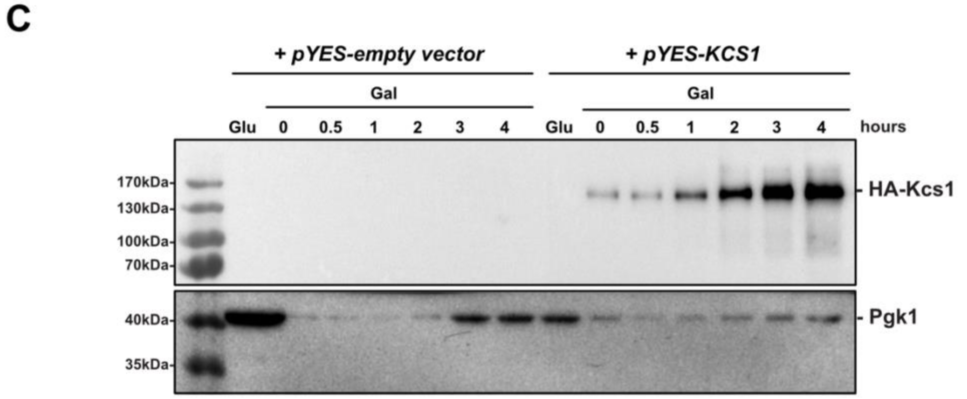


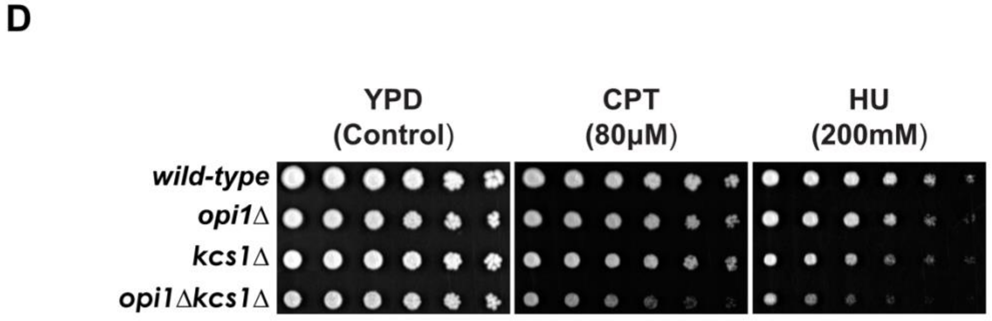

#### Supplementary Figure 9

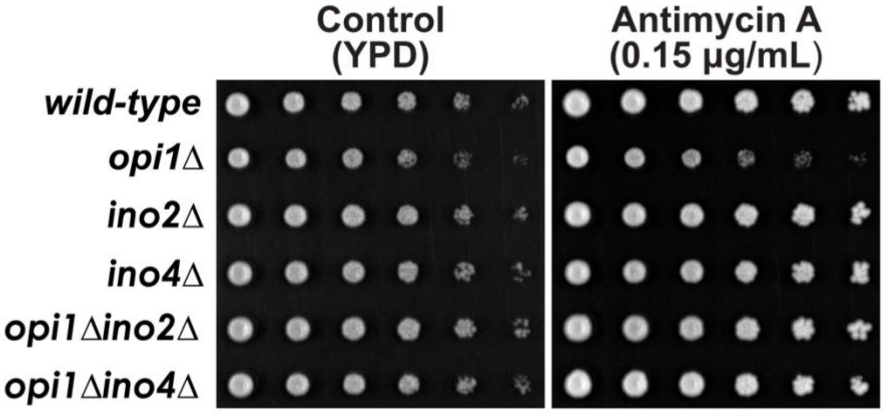

**Supporting Information**

**S1 Fig. Principal Component Analysis (PCA) plot showing the clustering of gene expression profiles in different experimental groups.**

The PCA plot is based on the first two principal components (PC1 and PC2) derived from RNA-seq data. Each data point represents an individual sample, and the colors indicate different experimental groups: MMS-treated *opi1Δ* (red), MMS-treated wild-type (green), untreated *opi1Δ* (blue), and untreated wild-type (purple).

**S2 Fig. The increased sensitivity of *opi1Δ* cells to MMS is not due to loss of viability.**

**(A)** Western-blot showing expression of Opi1-HA under the control of its endogenous promoter from a pRS416 plasmid.

**(B)** A pRS416 plasmid expressing Opi1-HA from its endogenous promoter rescues the MMS sensitivity of an *opi1Δ* strain.

**(C)** Colony forming unity assay (CFU) to assess cell viability after acute exposure to MMS. Exponentially growing wild-type and *opi1Δ* cells were subjected to acute exposure with 0.1% MMS for 60 minutes and 90 minutes in YPD medium. Following the treatment, cells were plated onto YPD agar plates in three technical replicates and incubated for 2-3 days at 30$^{o}$C to allow colony formation. Percentage of viable cells was determined based on the number of colonies formed on YPD plates from cells treated with MMS relative to control plates (untreated cells).

**(D)** Analysis of cell viability by flow citometry. Exponentially growing wild-type and *opi1Δ* cells were subjected to acute exposure with 0.1% MMS for 2 hours in YPD medium. t-tests were used to determine the statistical significance as explained in the Material and Methods section. Note: For (B) $p$ = 0.27 (wild-type MMS 60' vs *opi1Δ* MMS 60'); $p$ = 0.72 (wild-type MMS 90' vs *opi1Δ* MMS 90'). For (C) $p$ = 0.35 (wild-type vs *opi1Δ*).

**S3 Fig. Rad53 signaling is not affected in *opi1Δ* cells.**

Western-blot analysis shows the MMS-induced phospho-shift of Rad53 in the indicated strains. Asynchronous cultures in exponential growth phase were treated with genotoxins, and samples were collected at the specified time points for western-blot analysis using an antibody against Rad53.

**S4 Fig. Opi1-GFP translocates to the nucleus upon MMS treatment in the absence of inositol.**

Cellular localization of Opi1-GFP was scored manually from two independent experiments. For more details see Material and Methods section.

**S5 Fig. Opi1 is important to modulate gene expression during MMS-induced genotoxic stress.**

**(A)** Bar plot illustrating the top enriched biological processes identified in the gene set that are induced during MMS treatment in an Opi1-dependent manner (Table S6). Each bar represents a specific biological process, and the height of the bar corresponds to the fold enrichment with a p-adjusted value of 0.05. Gene ontology (GO) analysis was conducted in PANTHER using the overrepresentation test with *Saccharomyces cerevisiae* serving as the reference list, which includes all genes in database. The test type used was FISHER, and the false discovery rate (FDR) correction was applied.

**(B)** Gene Ontology (GO) term enrichment network depicting the relationships between genes grouped in clusters that are induced during MMS treatment in an Opi1-dependent manner (Table S6). The enrichment analysis was performed using the STRING database, which integrates various sources of functional information with a confidence score >0.7 (high confidence). Nodes in the network represent genes, while edges indicate the connections between them. The color of the nodes represents a biological process, and the thickness of the edges represents the increased confidence of the interaction between nodes.

**S6 Fig. Cellular response of Ino2 and Ino4-deficient cells to different stress signals.**

Serial dilution assays showing the effect of heat-shock, oxidative stress ($H_2O_2$), osmotic stress (KCl) and ER stress (tunicamycin) treatment upon the sensitivity of wild-type, *opi1Δ*, *ino2Δ*, *ino4Δ* and double mutants. Four-fold serial dilutions were spotted on YPD plates and grown for 2–3 days at 30°C.

**S7 Fig. Opi1 show negative genetic interactions with genes from phosphatidylinositol and phosphatidylcholine biosynthesis.**

**(A-B)** Deletion of genes from phosphatidylinositol and phosphatidylcholine synthesis synergistically increases the MMS sensitivity of *opi1Δ* cells.

**(C)** Cells overexpressing *INO1* under the control of the *GAL1* promoter can rescue the inositol auxotrophic of an *ino1Δ* strain.

**(D)** Overexpression of *INO1* does not phenocopy the MMS sensitivity of an *opi1Δ* strain. Fourfold serial dilutions were spotted on YPD (A-B) or SC -URA (C-D) with glucose or galactose and plates were grown for 2–3 days at 30°C.

**S8 Fig. Inositol pyrophosphates overproduction by Siw14 deletion and Kcs1 overexpression.**

**(A)** Cells lacking the inositol pyrophosphate phosphatase Siw14 show mild sensitivity to MMS**.**

**(B-C)** *KCS1* mRNA and protein levels were determined in cells overexpressing KCS1 from a pYES2-ntc plasmid under the control of the GAL1 promoter. Relative mRNA levels were quantified as fold change compared to a control condition (pYES-e/v 0'), where e/v represents empty vector. Real-time PCR was performed using an ABI Prism 7500 instrument (Applied Biosystems) with the Quantinova Probe PCR kit (Qiagen), utilizing TaqMan® probes specific for *KCS1* (Sc04136910_s1) and *ACT1* (Sc04120488_s1) as a reference gene (Thermo Fisher Scientific). For details on galactose induction and growth conditions, refer to the "Growth Conditions" section in the Material and Methods.

**(D)** An *opi1Δkcs1Δ* show sensitivity to replication stress inducers hydroxyurea (HU) and camptothecin (CPT).

For (A) and (D), fourfold serial dilutions were spotted on YPD plates and grown for 2–3 days at 30°C.

**S9 Fig. Deletion of Ino2-Ino4 and Kcs1 ameliorates mitochondria function in *opi1Δ* cells.**

**(A)** Deletion of Ino2 and Ino4 rescues the antimycin A sensitivity of *opi1Δ* cells.

**(B)** The *opi1Δkcs1Δ* double mutant exhibits restored mitochondrial respiratory capacity, which is impaired in the single *opi1Δ* and *kcs1Δ* mutants.

**S1 Table.** List of *S. cerevisiae* strains used in this study.

**S2 Table.** List of Plasmids used in this study.

**S3 Table**. List of primers used in this study.

**S4 Table.** Differentially expressed genes between the *opi1*Δ and wild-type strains in untreated condition ("genotype_OPI1_vs_WT.csv").

**S5 Table.** Differentially expressed genes in the wild-type strain treated with MMS relative to untreated ("treatment_MMS_vs_unt.csv").

**S6 Table.** Differential effect of MMS treatment in *opi1*Δ strain relative to the wild-type ("interaction.csv").

**S7 Table.** Differentially expressed genes in the *opi1*Δ strain treated with MMS relative to the *opi1*Δ without treatment ("treatment_MMS_vs_unt_OPI1.csv").

**S8 Table.** Differentially expressed genes between the *opi1*Δ and the wild-type strains both treated with MMS ("OPI1_vs_WT_treatment.csv").
